# Revealing the impact of recurrent and rare structural variants in multiple myeloma

**DOI:** 10.1101/2019.12.18.881086

**Authors:** Even H Rustad, Venkata D Yellapantula, Dominik Glodzik, Kylee H Maclachlan, Benjamin Diamond, Eileen M Boyle, Cody Ashby, Patrick Blaney, Gunes Gundem, Malin Hultcrantz, Daniel Leongamornlert, Nicos Angelopoulos, Daniel Auclair, Yanming Zhang, Ahmet Dogan, Niccolò Bolli, Elli Papaemmanuil, Kenneth C. Anderson, Philippe Moreau, Herve Avet-Loiseau, Nikhil Munshi, Jonathan Keats, Peter J Campbell, Gareth J Morgan, Ola Landgren, Francesco Maura

## Abstract

The landscape of structural variants (SVs) in multiple myeloma remains poorly understood. Here, we performed comprehensive classification and analysis of SVs in multiple myeloma, interrogating a large cohort of 762 patients with whole genome and RNA sequencing. We identified 100 SV hotspots involving 31 new candidate driver genes, including drug targets BCMA (*TNFRSF17*) and *SLAMF7*. Complex SVs, including chromothripsis and templated insertions, were present in 61 % of patients and frequently resulted in the simultaneous acquisition of multiple drivers. After accounting for all recurrent events, 63 % of SVs remained unexplained. Intriguingly, these rare SVs were associated with up to 7-fold enrichment for outlier gene expression, indicating that many rare driver SVs remain unrecognized and are likely important in the biology of individual tumors.

## Introduction

Structural variations (SVs) are increasingly recognized as key drivers of cancer development^1–3^. Functional implications of SVs are diverse, including large-scale gain and loss of chromosomal material, gene regulatory effects such as super-enhancer hijacking, and gene fusions^4^. The basic unit of SVs are pairs of breakpoints, classified as either deletion, tandem duplication, translocation or inversion^5^. Recently, whole genome sequencing (WGS) studies have revealed complex patterns where multiple SVs are acquired together in a single event, often involving multiple chromosomes, with a potentially important role in cancer initiation and progression^2,6–12^.

In multiple myeloma, chromosomal translocations are considered to be initiating events in ∼40 % of patients, where the immunoglobulin heavy chain (*IGH*) locus on chromosome 14 is juxtaposed with a class-defining oncogene^13^. Translocations involving *MYC* have been identified in 15-42 % of newly diagnosed multiple myeloma patients and are associated with progression to multiple myeloma from its precursor stage, smoldering multiple myeloma^14–17^. Recurrent translocations involving the immunoglobulin lambda (*IGL*) locus and other genes have recently been reported, expanding the catalogue of potential drivers and confirming the critical role of SVs in multiple myeloma pathogenesis^14^.

The genomic landscape of multiple myeloma is characterized by multiple aneuploidies, which fall into two main categories^13,18,19^. First, whole chromosome gains of more than two odd-number chromosomes is referred to as hyperdiploidy (HRD) and present in ∼60 % of patients^20^. HRD is considered to be the initiating event in most of the patients without a canonical *IGH*-translocation, with some patients acquiring additional whole chromosome gains later in tumor evolution^13,20^. The second category is made up of all other recurrent copy number alterations (CNAs), one or more of which are present in virtually all patients^19,20^. Among these, gain of chromosome 1q21 often occurs early in tumor evolution, while loss of tumor suppressor genes tends occur later^20–23^. The structural basis of these established drivers remains poorly understood, and there is a lack of data about the role of SVs beyond the most recurrent aberrations.

We recently reported the first comprehensive SV characterization using WGS of sequential samples from 30 multiple myeloma patients^20^. Despite the small sample set, SVs emerged as key drivers during all evolutionary phases of disease, and we identified a high prevalence of three main classes of complex SVs: chromothripsis, templated insertions and chromoplexy^20^. In chromothripsis, chromosomal shattering and random rejoining results in a pattern of tens to hundreds of breakpoints with oscillating copy number across one or more chromosomes^24^. Templated insertions are characterized by focal gains bounded by translocations, resulting in concatenation of amplified segments from two or more chromosomes into a continuous stretch of DNA, which is inserted back into any of the involved chromosomes^2,20^. Chromoplexy similarly connects segments from multiple chromosomes, but the local footprint is characterized by copy number loss^25^. Importantly, complex SVs represent large-scale genomic alterations acquired by the cancer cell at a single point in time, potentially driving subsequent tumor evolution.

Here, we present the first comprehensive study of SVs in a large series of 762 multiple myeloma patients. We show that recurrent and rare SVs are critical in shaping the genomic landscape of multiple myeloma, including complex events simultaneously causing multiple drivers.

## Results

### Genome-wide landscape of structural variation in multiple myeloma

To define the landscape of simple and complex SVs in multiple myeloma, we investigated 762 newly diagnosed patients from the CoMMpass study (NCT01454297) who underwent low coverage long-insert WGS (median 4-8X) (**Table S1**). RNA sequencing was also available from 592 patients. As a validation cohort, we compiled previously published WGS data from 52 patients^20,26,27^. For each patient sample, we integrated the genome-wide somatic copy number profile with SV data and assigned each pair of SV breakpoints as either simple or part of a complex event according to the three main classes previously identified in multiple myeloma (**Methods**)^20^. Templated insertions involving more than two chromosomes were considered complex. Events involving more than three breakpoint pairs which did not fulfill the criteria for a specific class of complex event were classified as unspecified “complex”^20^.

Overall, we identified a median of 15 SVs per patient in the CoMMpass cohort (interquartile range, IQR 7-28) (**Figure 1A**). Deletion was the most common SV type (33 %), followed by inversion and translocation (24 % each), and tandem duplication (18 %) (**Figure S1A**). Fifty-one percent of SVs (8842/17185) were defined as part of 931 complex events (**Figure 1A**). Chromothripsis, chromoplexy and templated insertions involving >2 chromosomes were observed in 21%, 11% and 21 % of patients, respectively; 32 % of patients had an unspecified complex event. One or more complex events were identified in 61 % of patients (median 1; range 0-14); 16 % had two complex SVs; 14 % had three or more. The distribution across simple and complex classes varied for each SV type: deletions were most commonly simple events (71 %), followed by 18 % attributed to chromothripsis; while translocations were simple events in 28 % (e.g. reciprocal or unbalanced) and part of templated insertions or chromothripsis in 31 % each (**Figure S1A**). In the validation cohort we identified a slightly higher overall SV burden, with a median of 26 SVs per patient (IQR 10-36), as expected from the higher sequencing coverage. Reassuringly, the distribution across simple and complex SV classes was similar across the datasets (**Figure S1A**), suggesting that low-coverage long-insert WGS provides a representative view of the SV landscape.

**Figure 1:**
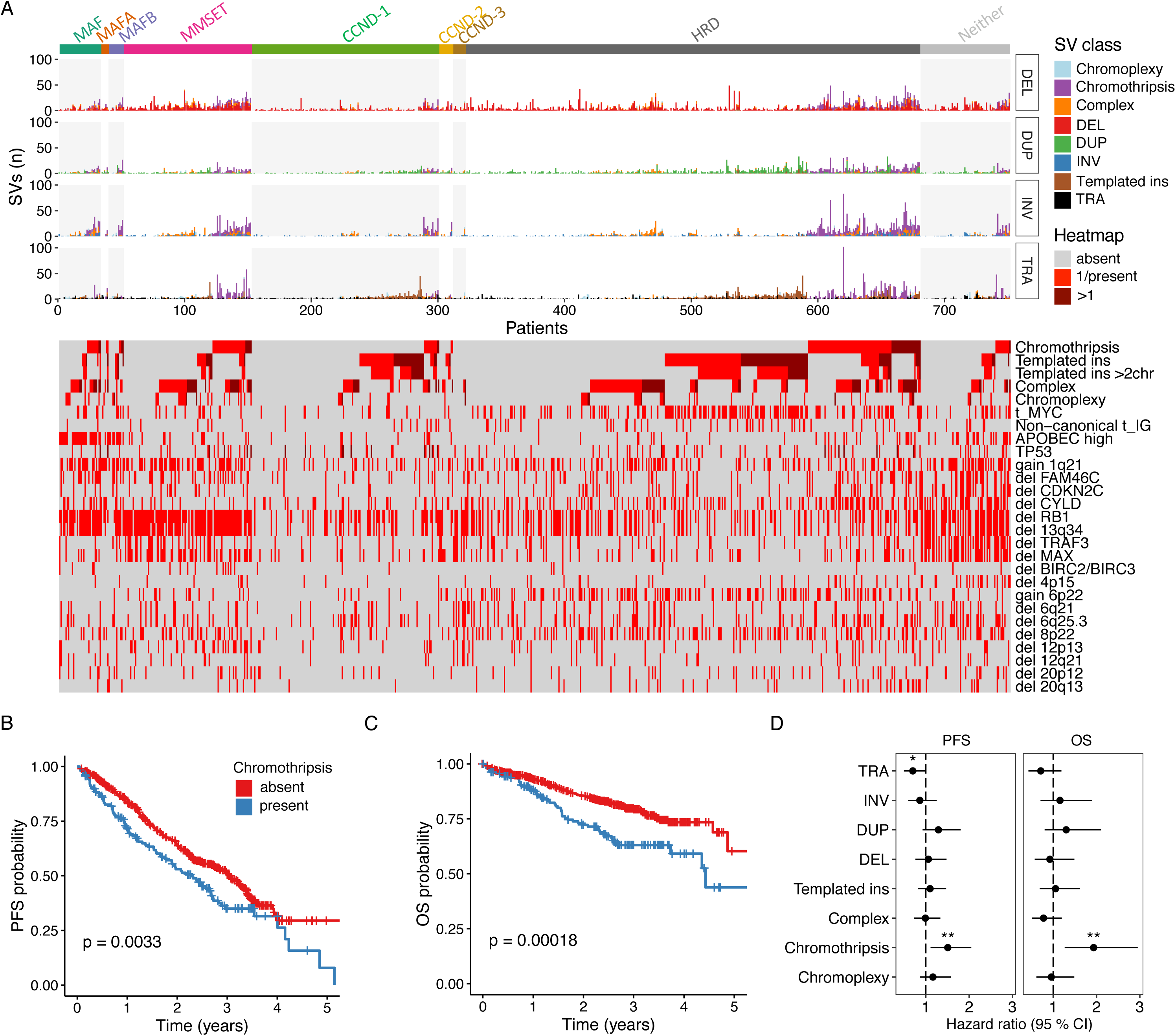
Simple and complex classes of structural variants in multiple myeloma. **A:** Genome-wide burden of structural variant types (horizontal panels) in each patient, grouped according to primary multiple myeloma molecular subgroup. Bars within each panel are colored according to the contribution of complex SV classes (i.e. chromoplexy, chromothripsis, templated insertions or other ‘complex’ variants grouped together) versus simple SVs (DEL, deletion; DUP, tandem duplication; INV, inversion and TRA, translocation). Heatmap below shows the presence of complex structural variant classes together with key genomic drivers of multiple myeloma, including non-canonical immunoglobulin (IG) translocations. **B-C:** Kaplan-meier plots for progression free survival (PFS) **(B)** and overall survival (OS) **(C)** in patients with and without chromothripsis (shown in blue and red respectively). **D**: Hazard ratio for PFS and OS by SV type, estimated using multivariate Cox regression. (Line indicates 95 % CI from multivariate cox regression models, statistically significant features indicated by asterisks (* p<0.05; ** p < 0.01). For templated insertions, chromothripsis, chromoplexy and unspecified complex events, we compared patients with 0 versus 1 or more events. The remaining SVs were considered by their simple class (i.e. DUP, DEL, TRA and INV), comparing the 4^th^ quartile SV burden with the lower three quartiles. The multivariate models included all SV variables as well the following clinical and molecular features: age, sex, ECOG status, ISS-stage, induction regimen, gain1q21, delFAM46C, delTRAF3, delTP53, delRB1, high APOBEC mutational burden, hyperdiploidy and canonical translocations involving *CCND1*, *MMSET*, *MAF*, *MAFA*, *MAFB* and *MYC*.

SV breakpoints tend to be enriched within specific genomic contexts, depending on the underlying mechanism. Our findings from multiple myeloma corresponded with previous reports from pan-cancer studies^2,11^. A higher density of SV breakpoints was generally associated with early replication, chromatin accessibility (DNAse seq), highly expressed genes and active enhancer regions as defined by histone H3K27 acetylation (H3K27ac) (**Figure S1B; Methods**). These associations were particularly strong for the breakpoints of tandem duplications and templated insertions, which also showed a strong correlation with each other (Spearman’s rho = 0.33, p < 0.001). Turning to the SV burden in individual patients, the number of templated insertion breakpoints was inversely correlated with all other SV classes, except tandem duplications, which showed a striking positive correlation (Spearman’s rho = 0.46, p <0.001) (**Figure S1C**). Taken together, templated insertions and tandem duplications tended to occur in the same genomic contexts, and to be enriched in the same tumors.

In patients with newly diagnosed multiple myeloma, complex SV classes showed distinct patterns of co-occurrence, mutual exclusivity and association with recurrent molecular alterations (**Figures 1A** **and S2**). High APOBEC mutational burden, a feature associated with aggressive disease and poor outcomes^28,29^, was enriched in patients with chromothripsis (OR = 2.8, 95% CI 1.8-4.3; p < 0.001) and unspecified complex events (OR = 2.6, 95% CI 1.7-4.0; p < 0.001), but was mutually exclusive with templated insertions (OR = 0.49, 95% CI 0.29-0.80; p = 0.003). Chromothripsis was the only SV class with clear prognostic implications after adjustment for molecular and clinical features, resulting in adverse progression free survival (PFS, adjusted HR = 1.57; 95% CI 1.13-2.22; p = 0.008) and overall survival (OS, adjusted HR = 2.4; 95% CI 1.5-3.83; p < 0.001) (**Figure 1B-D; Methods**)^30^.

Bi-allelic inactivation of *TP53* was identified in 23 patients in our cohort, defining a patient population with very poor prognosis^21^. Interestingly, these patients showed strong enrichment for chromothripsis (15 out of 24; OR 7.8, p < 0.001) and high APOBEC mutational burden (OR 6.6, p < 0.001) (**Figure 1A** and **S2**). In 3 of 15 cases, the *TP53* deletion was caused by a chromothripsis event; and the association remained highly significant after excluding these cases (OR 6.2, p < 0.001). This relationship did not reach statistical significance for monoallelic *TP53* events. Consistent with our findings, an association between chromothripsis and TP53 inactivation has been reported in several other cancers^31–33^.

### Structural basis of recurrent translocations and copy number variants

To define the structural basis of canonical translocations in multiple myeloma, we identified all translocation-type events (single and complex) with one or more breakpoints involving the immunoglobulin loci (i.e. *IGH*, *IGK* and *IGL*) or canonical *IGH*-partners (e.g. *CCND1*, *MMSET* and *MYC*)^13^. Templated insertions emerged as the cause of *CCND1* and *MYC* translocations in 34 % and 73 % of cases, respectively (**Figure 2A**). This is particularly important given that templated insertions connect and amplify distant genomic segments, often involving several oncogenes and regulatory regions (e.g. super-enhancers). Indeed, templated insertions of *CCND1* and *MYC* were associated with focal amplification in 63 % and 90 % of cases, respectively; and involved more than two chromosomes in 24 % and 41 % of cases. Although rare, we also found examples of chromothripsis and chromoplexy underlying canonical *IGH* translocations, resulting in overexpression of the partner gene consistent with a driver event (**Figure 2B**). Twenty-four patients (3.2%) had a translocation involving an immunoglobulin locus (*IGH* = 11, *IGL* = 12 and *IGK* = 1) and a non-canonical oncogene partner, most of which are key regulators of B-cell development (e.g. *PAX5* and *CD40*) (**Figure 2B**)^34,35^. One of the patients with non-canonical *IGH* translocation also had an *IGH*-*MMSET* translocation; while the remaining 10 patients either had HRD (n = 8) or no known initiating event (n=2), suggesting the disease may have been initiated by a non-canonical *IGH* translocation. Taken together, we show that different mechanisms of SV converge to aberrantly activate key driver genes in multiple myeloma. Importantly, SVs with strong biological implications and a potential role in tumor initiation are not necessarily recurrent, even in a series as large as the one analyzed here.

**Figure 2:**
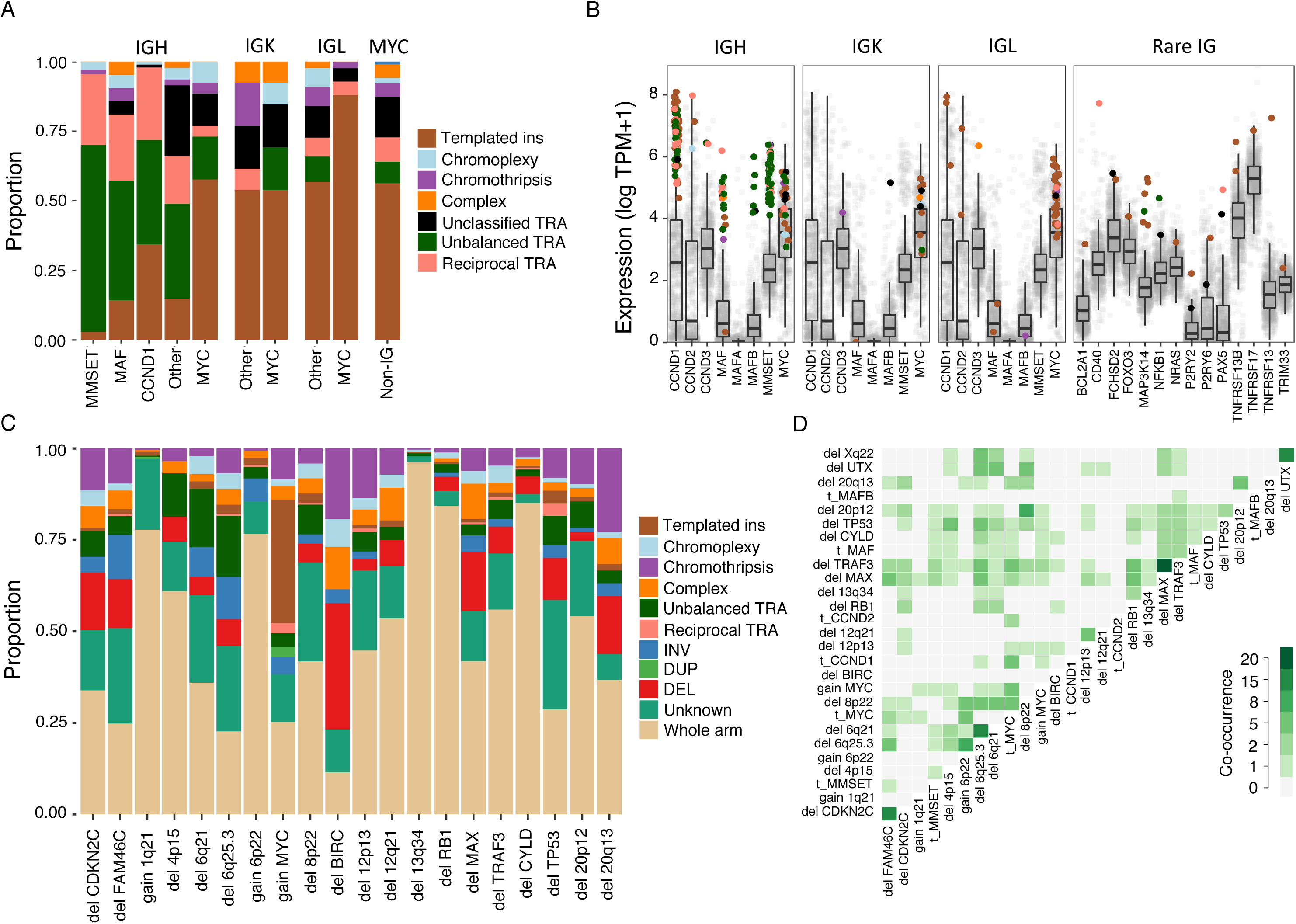
Structural basis of recurrent translocations and copy number changes. **A:** Relative contribution (y-axis) of simple and complex SV classes to translocations (TRA) involving the immunoglobulin loci and *MYC* with canonical and non-canonical partners (x-axis). The category “Other” contains non-*MYC* translocations for *IGK* and *IGL*, while for *IGH* it includes *CCND2*, *CCND3*, *MAFA* and *MAFB*. **B:** Transcript per million (TPM) gene expression of canonical and non-canonical partners of translocations involving the immunoglobulin (IG) loci. Each point represents a sample, colored by the SV class involved or absence of SV (gray). Boxplots shows the median and interquartile range (IQR) of expression across all patients, with whiskers extending to 1.5 * IQR. The templated insertion of *IGH* and *MAF* with low expression was part of a multi-chromosomal event involving and causing the overexpression of *CCND1*. **C)** Structural basis of recurrent multiple myeloma CNA drivers, showing the relative contribution of whole arm events and CNAs associated with a specific SV. Intrachromosomal events without a clear causal SV were classified as “unknown”. **D)** Co-occurrence matrix showing all cases where two or more recurrent CNAs and/or canonical driver translocations were caused by the same simple or complex SV. Each CNA segment was only counted once, even if including more than one driver gene. For example, a deletion involving both *CDKN2C* and *FAM46C* will be reported only if connected to another CNA caused by the same SV. BIRC: *BIRC2*/*BIRC3*.

Next, we addressed the structural basis of known recurrent CNAs (**Table S2**). Aneuploidies involving a whole chromosome arm were most common (60 % of 2926 events); 24 % could be attributed to a specific SV and 14 % were intrachromosomal events without a clear causal SV, classified as “unknown” (**Figure 2C**). The proportion of CNAs explained by a SV was similar in the validation dataset (29 % of 450 events; Fisher’s test p= 0.41; **Table S3**). There was considerable variation in the proportion and class of SVs causing gains and losses between different loci, indicating the presence of distinct underlying mechanisms being active at these sites (**Figure 2C**). Loss of *CDKN2C*, *FAM46C*, 6q21, 6q25.3, *MAX* and *TP53* could be attributed to SVs in ∼50 % of cases. SVs accounted for 78 % of *BIRC2*/*BIRC3* losses, and 62 % of *MYC* gains, of which 34 % were caused by templated insertions. Interestingly, unbalanced translocations were responsible for 5.5 % of all recurrent deletions (n = 132), up to 8 % for 17p and 16 % for 6q. Finally, 44.5 % of all chromothripsis events resulted in the acquisition of at least one recurrent driver CNA (n = 94); the corresponding numbers for chromoplexy and templated insertions involving >2 chromosomes were 41.6 % (n = 40) and 19.9 % (n = 42), respectively.

In 12.5% of patients (n=95), two or more seemingly independent recurrent CNAs or canonical translocations were caused by the same simple or complex SV (**Figure 2D**). The most common event classes were chromothripsis (n = 33), chromoplexy (n = 22), unbalanced translocation (n = 21) and templated insertion (n = 21) (**Figure 3A-C**). Intriguingly, translocation of *MYC* occurred as part of the initiating *IGH*-translocation in five patients, together with *CCND1* (n=3) or *CCND2* (n=2). Another interesting phenomenon was observed on chromosome 6, where gain of one arm (p) and loss of the other (q) resulted from a single inversion event in seven patients (**Figure 3D**).

**Figure 3:**
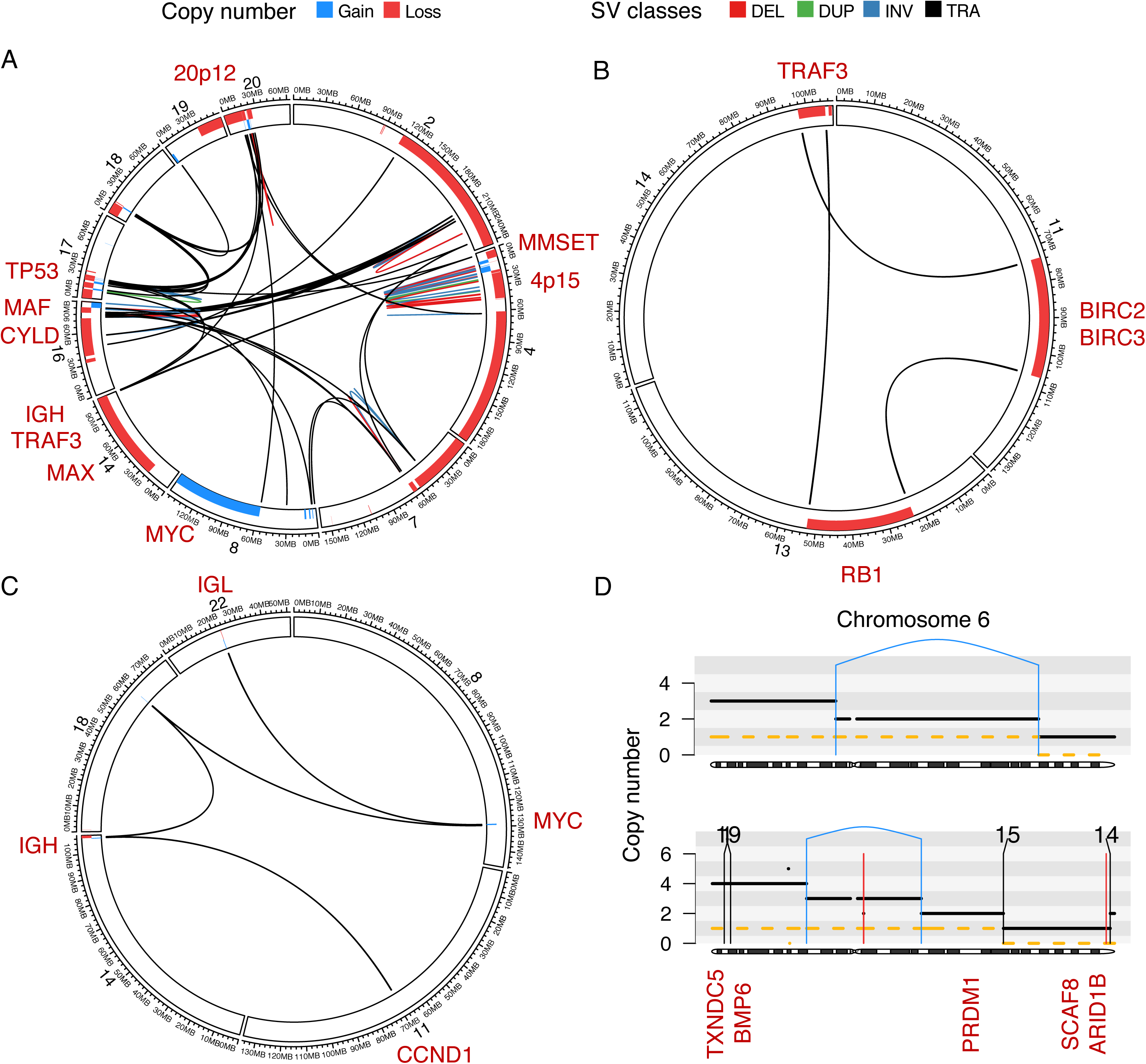
Multiple recurrent aberrations caused by the same SV. **A-C)** Genome plots showing the SV breakpoints of a single complex SV in each panel (colored lines), with bars around the plot circumference indicating copy number changes. The outer track represents copy number loss (red), the inner represents copy number gains (blue). **A)** Chromothripsis involving *IGH* and 9 recurrent driver aberrations across 10 different chromosomes (sample MMRF_1890_1_BM). **B)** Chromoplexy involving chromosomes 11, 13, and 14, simultaneously causing deletion of key tumor suppressor genes on each chromosome (sample MMRF_2194_1_BM). **C)** Templated insertion involving 5 different chromosomes, causing a canonical *IGH-CCND1* translocation and involving *MYC* in the same chain of events. Focal amplifications can be seen corresponding to the translocation breakpoints (sample MMRF_1862_1_BM). **D)** Top: Unbalanced inversion on chromosome 6, simultaneously causing gain of 6p and loss of 6q (sample MMRF_1740_1_BM). Bottom: Copy number profile of chromosome 6 in sample MMRF_2266_1_BM, generated through several independent structural variants. The main events were: 1) whole chromosome gain; 2) unbalanced inversion causing an additional gain of 6p and loss of one of the two duplicated 6q alleles; and 3) chromoplexy involving chromosome 14 and 15, with loss of an interstitial segment containing *SCAF8* and *ARID1B* on 6q. This patient also had an independent chained translocation event involving the *TXNDC5*/*BMP6* locus on 6p.

### Hotspots of structural variation

Twenty recurrently translocated regions have been previously reported in multiple myeloma, defined by a translocation prevalence of >2 % within 1 Mb bins across the genome^14^. Recurrent regions included the immunoglobulin loci, canonical partner oncogenes, as well recurrent *MYC* partners, such as *BMP6*/*TXNDC5*, *FOXO3* and *FAM46C*^14,17,36^. We were motivated to expand the known catalogue of genomic loci where SVs play a driver role in multiple myeloma and are therefore positively selected (i.e. SV hotspots), considering all classes of single and complex SVs. To accomplish this, we applied the Piecewise Constant Fitting (PCF) algorithm, comparing the local SV breakpoint density to an empirical background model (**Methods; Figure S3A-E; Data S1**)^8,37^. Overall, we identified 100 SV hotspots after excluding the immunoglobulin loci (i.e., *IGH*, *IGL* and *IGK*), 85 of which have not been previously reported (**Figure 4A**; **Table S4**). Each patient had a median of three hotspots involved by a SV (IQR 1-5); and the number of hotspots involved was strongly associated with the overall SV burden (Spearman’s Rho = 0.51; p < 0.001; **Figure S3F**). Two of the previously reported regions of recurrent translocation were not recapitulated by our hotspot analysis: a locus of unknown significance on 19p13.3, and the known oncogene *MAFB* on 20q12. This may be explained by the behavior of the PCF algorithm, which favors the identification of loci where breakpoints are tightly clustered compared with neighboring regions as well as the expected background. Translocations involving *MAFB* were identified in 1.6 % of patients in our series (n = 12), but the breakpoints were widely spread out across 600 Kb in the intergenic region upstream of *MAFB*^38^. Similarly, a 2 Mb region on 19p13.3 was involved in 44 patients (5.7 %), predominantly by templated insertions and tandem duplications, but the breakpoints did not form a distinct cluster.

**Figure 4:**
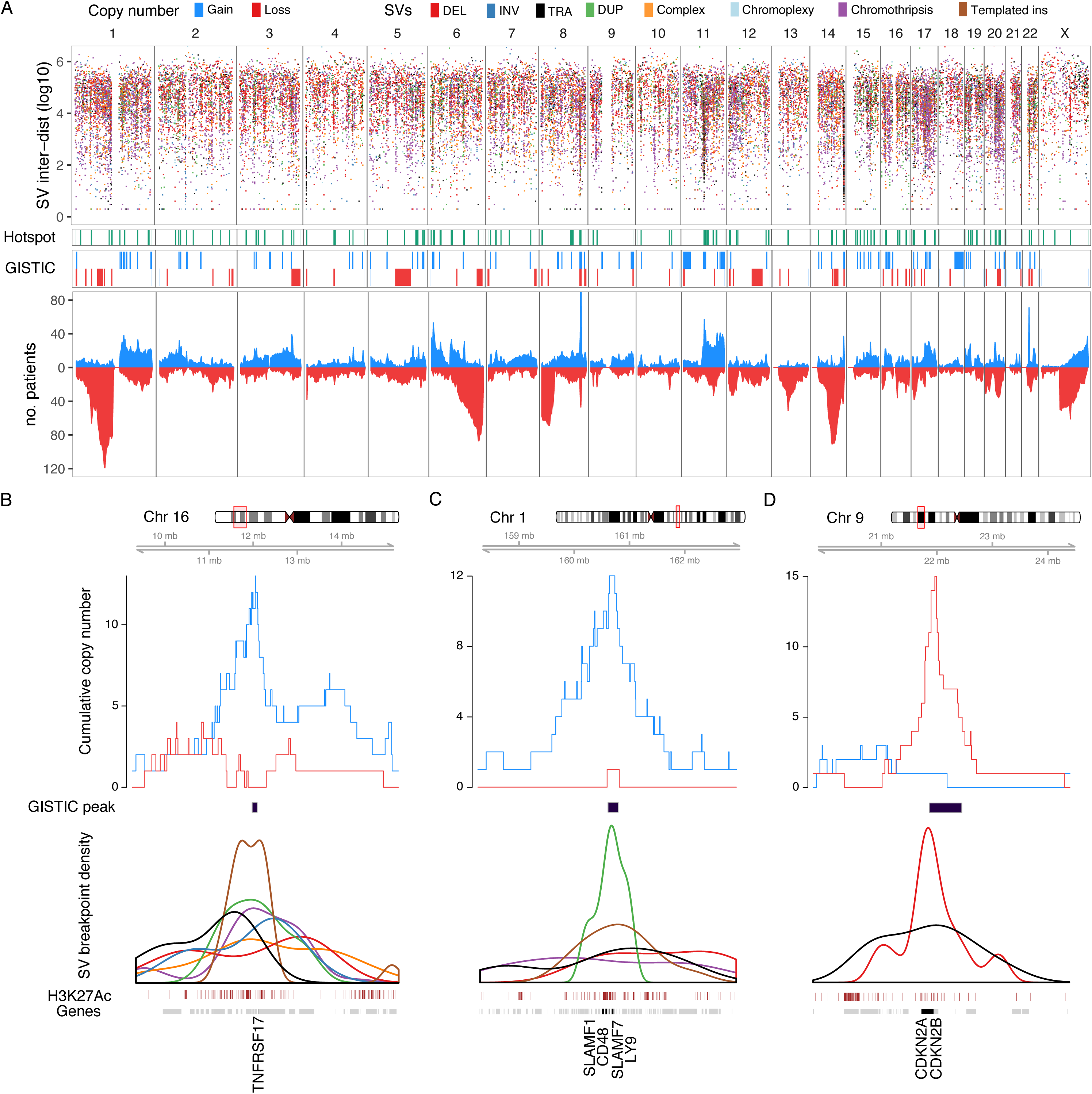
Genome-wide distribution of structural variation breakpoints and hotspots. **A)** Top: Rainfall plot showing the log-distance of each SV breakpoint (points) to its closest neighbor (y-axis) across the genome (x-axis), colored by SV class (legend above figure). Middle: Distribution of SV hotspots (green) and recurrent copy number changes (red/blue) identified by the GISTIC algorithm. Bottom: all copy number changes caused by SV breakpoints, showing cumulative plots for gains (blue) and losses (red). **B-D)** Zooming in on three SV hotspots, showing the chromosomal region (top), cumulative copy number and GISTIC peaks (middle), and the breakpoint density of relevant SV classes (colors indicated in legend above **A**) around the hotspot (bottom). The lower plots are annotated with enhancer marks (histone H3K27ac) in brown and genes in gray, with putative driver gene names in black. **B)** Gain of function hotspot centered around *TNFRSF17* (BCMA), dominated by highly clustered templated insertions, associated with focal copy number gain of *TNFRSF17*. **C)** Gain of function hotspot involving four genes in the Signaling Lymphocyte Activation Molecule (SLAM) family of immunomodulatory receptors, including the gene encoding the monoclonal antibody target *SLAMF7*. The SV hotspot region was supported by a GISTIC peak of copy number gain. **D)** Deletion hotspot associated with copy number loss centered on the cyclin dependent kinase inhibitors *CDKN2A*/*CDKN2B*, supported by a peak of copy number loss identified by GISTIC.

Given that SVs and CNAs reflect the same genomic events, we hypothesized that functionally important SV hotspots would be associated with a cluster of CNAs. We therefore performed independent discovery of driver CNAs using GISTIC (genomic identification of significant targets in cancer)^39^. This algorithm identifies peaks of copy number gain or loss which are likely to contain driver genes and/or regulatory elements based on the frequency and amplitude of observed CNAs (**Figure 4A, Table S5-6**). In addition, we generated cumulative copy number profiles for the patients involved by SV at each hotspot. Finally, we evaluated the impact of SV hotspots on the expression of nearby genes. By integrating SV, CNA, and expression data, we went on to determine the most likely consequence of each hotspot in terms of gain-of-function, loss-of-function, and potential involvement of driver genes and regulatory elements (**Figures 4A-D** and **S4**).

Gain-of-function hotspots were defined by copy number gains and/or overexpression of a putative driver gene (**Table S4** and **Figures 4A-C**, **5A** and **S4**). Hotspots defined by these criteria showed overwhelming dominance of tandem duplication, translocation and templated insertion SV breakpoints (median 73 % of breakpoints [IQR 63-87 %] in gain-of-function hotspots vs 21 % [11-28 %] in other hotspots; Wilcoxon rank sum test p < 0.001; **Figure 5B**). By contrast, chromothripsis, chromoplexy and unspecified complex events had a similar contribution to gain- and loss-of-function hotspots (Wilcoxon rank sum test, Bonferroni-Holm adjusted p-values > 0.05). Strikingly, gain-of-function hotspots showed 5.5-fold enrichment of super-enhancers as compared with the remaining mappable genome (1.6 vs 0.3 super-enhancers per Mb, Poisson test p < 0.001)(**Methods**)^40^. Enhancer peaks (H3K27ac) clustered around the SV breakpoints were observed in 46 out of 62 gain-of-function hotspots (74 %) (**Figure 5A**). High-confidence driver genes supported by both overexpression and an amplification peak: *CCND1*, *CCND2*, *MYC*, *NSD2* (*WHSC1)*, *P2RY2/FCHSD2*, *FOXO3*, *MAP3K14*, *CD40*, *LTBR*, *TNFRSF17*, *PRKCD*, *SEC62*/*SAMD7*, *SOX30* and *GLI3* (**Figure 5A**). Of particular interest, *TNFRSF17* was involved by SVs in 5 % of patients (n = 38) and encodes B-Cell Maturation Antigen (BCMA), a therapeutic target of chimeric antigen receptor T-cells (CAR-T) and monoclonal antibodies (**Figure 4B**)^41^. We also confirmed previous reports that virtually all SVs involving *MYC* resulted in its overexpression, including deletions and inversions, which acted by repositioning *MYC* next to the super-enhancers of *NSMCE2*, roughly 2 Mb upstream^42^. Numerous genes were involved by gain-of-function hotspots without clear evidence of overexpression, many of which are known to be important in multiple myeloma, such as *IRF4*, *KLF2, KLF13* and *SLAMF7* (**Table S4** and **Figure 4C**). Overall, translocation-type SVs involving an immunoglobulin locus (i.e., *IGH*, *IGL* or *IGK*), either directly or as part of a complex event, were associated with the strongest gene expression effects (pairwise Wilcoxon rank sum test, Bonferroni-Holm adjusted p-values < 0.0001) (**Figure S5**). This was true both for canonical driver genes (median Z-score 1.7, IQR 1.1-2.4) and other putative drivers (median Z-score 1.83, IQR 0.85-2.5). Non-immunoglobulin translocations had the second highest impact, followed by tandem duplications and other SVs, all of which were associated with significantly higher expression than patients without SV (adjusted p-values < 0.01) (**Figure S5**).

**Figure 5:**
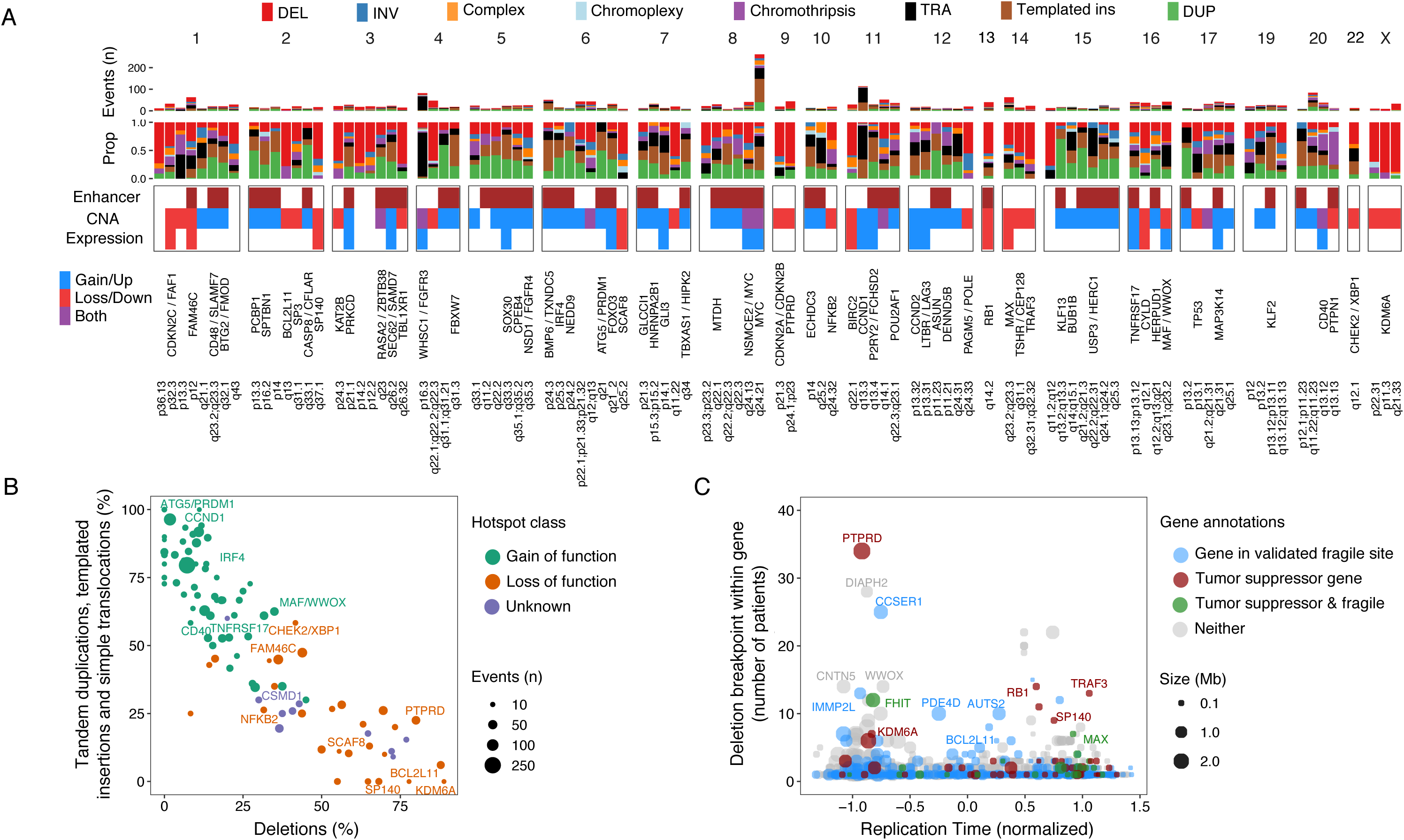
Gain- and loss-of function consequences of structural variant hotspots. **A)** Summary of all 100 SV hotspots, showing (from the top): absolute and relative contribution of SV classes within 100 Kb of the hotspot; involvement of active enhancers in multiple myeloma, copy number changes and differential expression of putative driver genes; known and candidate driver genes. **B**) Distribution of SV classes involving each hotspot, colored by gain-of-function (green), loss-of-function (orange) or unknown functional status (blue) as defined by copy number and gene expression data. Percentage of templated insertion, simple translocation and tandem duplication SVs on the y-axis and deletion SVs x-axis. **C**) Potential fragile sites in multiple myeloma, characterized by deletions (y-axis) within the body of large genes (size) that are late replicating (x-axis). Fragile sites indicated by the color scale have been validated in pan-cancer studies.

Loss-of-function hotspots, defined by copy number loss and/or reduced expression of a putative driver gene, were highly enriched for deletion breakpoints compared to the remaining hotspots (median 55 [IQR 40-68] % vs 13 [7-23] %; Wilcoxon rank sum test p < 0.001; **Table S4** and **Figures 4D**, **5A-B** and **S4**). High-confidence drivers defined by copy number loss and significantly reduced gene expression were *FAM46C*, *FAF1*, *RB1*, *MAX*, *SP140L*, *SCAF8*, *BIRC2* and *CYLD* (**Figure 5A**). Additional genes identified to be associated with loss of copy number, but without a clear effect on expression, included established tumor suppressor genes such as *TP53* and *CDKN2C,* as well as novel candidate drivers, e.g., *SP3* and *BCL2L11* (**Table S4**). Some genes with particular enrichment of intragenic deletion breakpoints were large and late-replicating, suggesting they may be fragile sites in multiple myeloma, prone to breaking during replication stress (**Figure 5C; Table S4** and **S7**)^43^. The top fragile gene candidate in our data was *PTPRD*, which is suspected to be a tumor suppressor gene in several cancers (**Figure 5C**)^27,43,44^. Additional large genes involved by recurrent intragenic deletions in this study included *CCSER1*, *FHIT* and *WWOX*, all of which are known fragile sites in cancer^43,45–47^. However, in a recent pan-cancer study, the accumulation of complex nested deletion SVs (termed *rigma*) in fragile site-associated genes including *FHIT* and *WWOX* could not be fully explained by the classical features of fragility, indicating that they may be positively selected^6^. Careful validation of the role played by these hotspots will be necessary.

Consistent with recent findings in breast cancer^37^, in our series, SV hotspots showed seven-fold enrichment for multiple myeloma germline predisposition SNPs as compared with the remaining mappable genome (6 SNPs in 125 Mb of hotspots vs 17 in the remaining genome, Poisson test p = 0.0002; **Figure S6**)^48,49^. The involved SNPs were on 3q26.2 (SV hotspot involving *SEC62/SAMD7*), 6p21.3 (unknown SV target), 6q21 (*ATG5/PRDM1*), 9p21.3 (*CDKN2A/CDKN2B*), 19p13.11 (*KLF2*) and 2q31.1 (*SP3*) (**Table S8**). One additional SNP on 8q24.21 (*CCAT1*) was less than 1 Mb upstream of *MYC*, between two SV hotspots centered around *NSMCE2* and *MYC,* respectively. Assuming that multiple myeloma predisposition SNPs and driver SVs affecting the same locus would have similar consequences, this co-occurrence may provide novel insights into the target genes and functional implications.

In our validation dataset of 52 genomes, 35 out of 100 hotspots showed significant enrichment of SV breakpoints compared with the remaining mappable genome (Poisson test, FDR < 0.1; **Table S9 and Figure S7A**). Hotspots that were confirmed in the validation dataset showed significantly higher prevalence in CoMMpass as compared with hotspots that were not confirmed (median 4 [IQR 2.6-5.4] % vs 2.2 [1.5-3] %; Wilcoxon rank sum test p<0.001).

Taken together, we identified 62 gain of function and 28 loss of function hotspots; with 10 hotspots being of unknown function (**Figures 5A, Table S4**). Thirty-three hotspots involved genes with established tumor suppressor or oncogene function in multiple myeloma; 31 additional hotspots involved novel putative driver genes (**Table S4**; all novel putative driver hotspots are shown in Figures 4B-C and **S4**).^50^

### Templated insertions are highly clustered and often involve multiple hotspots

Templated insertion breakpoints were highly clustered across the genome (**Figure 6A**) and associated with copy number gain in 76 % of cases (95 % CI 74-78 %), but only rarely with copy number loss (8.3 %; 95 % CI 7.0-9.6 %). Gains were almost exclusively single copy (92.5 % of 1200 gains), with less than one percent involving three or more copies above the baseline (n = 9), highlighting the stability of these events. Strikingly, out of 449 templated insertion events, 73 % involved one or more SV hotspots (including the immunoglobulin loci) (**Figure 6B-C**). Half of the templated insertions (n = 210) involved more than two chromosomes and 45 (10 %) involved more than four chromosomes.

**Figure 6:**
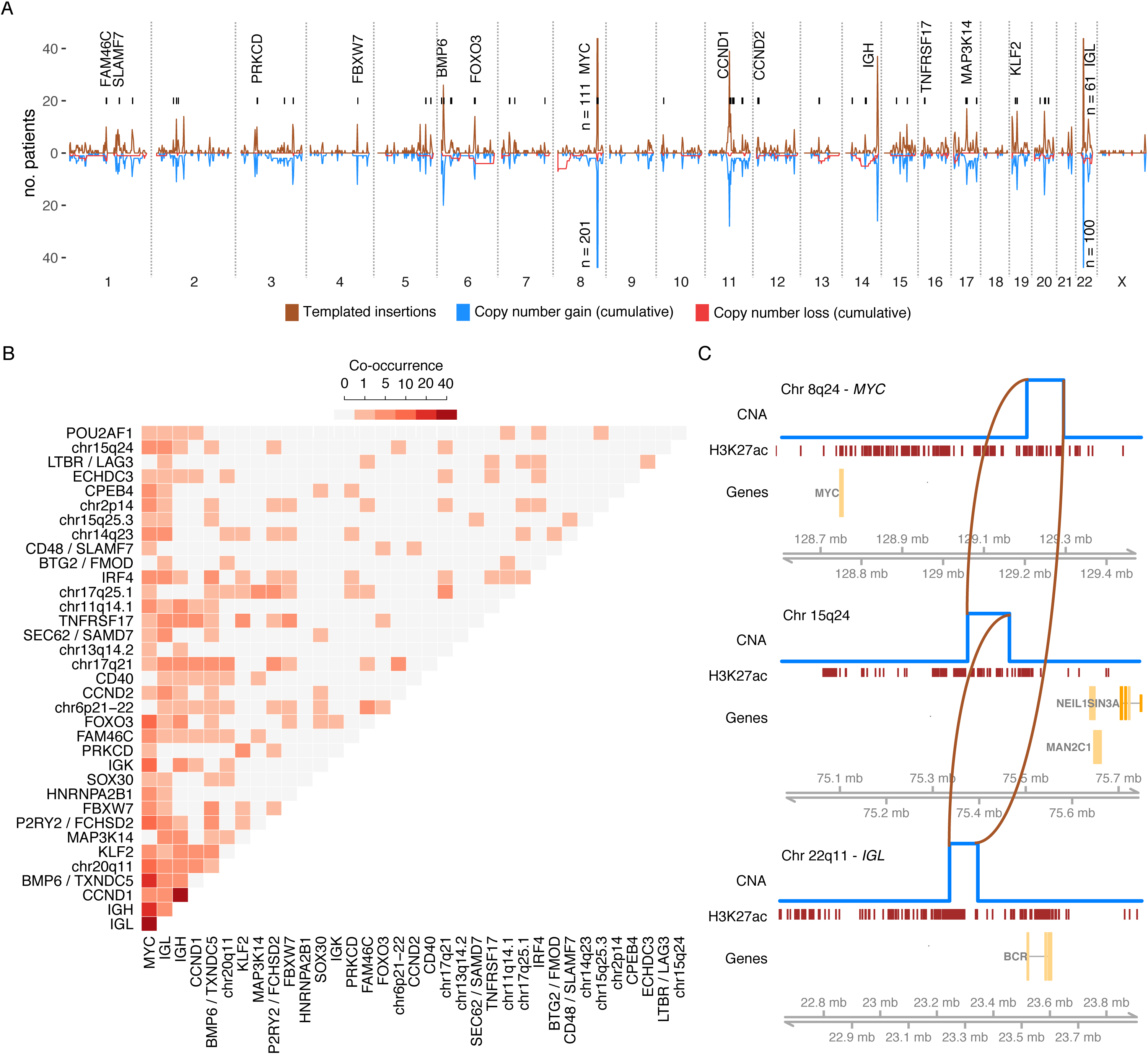
Templates insertions are highly clustered and string together multiple hotspots. **A)** Distribution of templated insertions across the genome (above) and associated cumulative copy number changes (below). Vertical black lines indicate SV hotspots containing >3 patients with a templated insertion. Selected gene names are annotated. Numbers are annotated where peaks extend outside of the plotting area **B)** Co-occurrence matrix showing the number of cases where pairs of SV hotspots occur as part of the same templated insertion, involving two or more chromosomes. **C)** Example of a templated insertion chain (brown lines) spanning three SV hotspots, involving the *IGL*, *MYC*, and a hotspot on chromosome 15q24 (sample MMRF_1550_1_BM). Copy number profile shown in blue, with an active enhancer track below (H3K27Ac).

Among the 210 events spanning more than two chromosomes, 39 % involved *MYC* or *CCND1*; 34 % involved *IGH, IGL* or *IGK*; and 30 % involved none of the above, but one or more other hotspots (**Figure 6C**). Overall, 54 % of multi-chromosomal templated insertions involved two or more hotspots (**Figure 6B-C**). In summary, we show that templated insertions in multiple myeloma are highly clustered and often string together amplified segments from multiple known drivers and SV hotspots in a single event.

### Chromothripsis is associated with wide-spread gain and loss of gene function mechanisms

In contrast to templated insertions, breakpoints of chromothripsis did not show strong focal clustering (**Figure 7A**). Instead, running the PCF algorithm for chromothripsis revealed enrichment in wider regions (**Table S10**). (**Figure 7A**). The most commonly involved loci included a 32 Mb region on chromosome 20q11-13 (n = 22 patients involved); 40 Mb spanning the *MYC* locus on 8q22-24 (n = 18); 23 Mb on 17q12-23 (n = 16); and 12 Mb on 12p12-13 (n=16). An important feature of chromothripsis is its ability to cause both gain- and loss-of-function as part of the same event^51^. Indeed, the breakpoints of chromothripsis were associated with chromosomal loss in 50.1 % of cases (95 % CI 48.8-52.5%) and gain in 39.6 % (95 % CI 37.7-41.5 %). High-level focal copy number gains (>5 copies) were predominantly caused by chromothripsis (75 % of segments) and present in only 6 % of patients overall (n = 46) (**Figure 7B-C**). In contrast to solid tumors, where chromothripsis may result in double minute chromosomes with >50 copies^2,6,10,52^, we observed no segments with more than 9 copies in this series. Similarly, in the validation cohort, the highest copy number associated with chromothripsis was 7 (**Figure S7B**). Genes involved by high-level focal gains had a two standard deviations higher expression, on average, as compared with the same genes in patients with normal copy number (median Z-score 2.2 vs −0.12, Wilcoxon rank sum test, p < 0.0001) (**Figure 7D**). Interestingly, despite the profound overall impact on gene expression, only four events (2.4 %) caused amplification and overexpression (Z-score > 2) of one or more established or putative drivers in multiple myeloma (i.e., *CCND1*, *CD40*, *NFKB1*, *MAP3K14* and *TNFRSF17*)^13^. For example, one patient (MMRF_2330) had amplification and overexpression of *MAP3K14* by chromothripsis, along with multiple genes that have not been reported as genomic drivers in multiple myeloma (**Figure 7B**). This included amplification and overexpression of the oncogene *RAD51C*, which is commonly amplified in breast cancer and may play a drive role in this one patient with multiple myeloma^53^. Two additional patients had >6 copies and outlier overexpression of *RPS6KB1*, encoding the 70 kDa ribosomal protein S6 kinase (p70S6K), a crucial regulator of cell cycle, growth and survival in the PI3K/mTOR pathways (**Figure S7C**)^54^. These observations raise questions regarding the impact of rare SVs in deregulating potential drivers, and their contribution to multiple myeloma biology overall.

**Figure 7:**
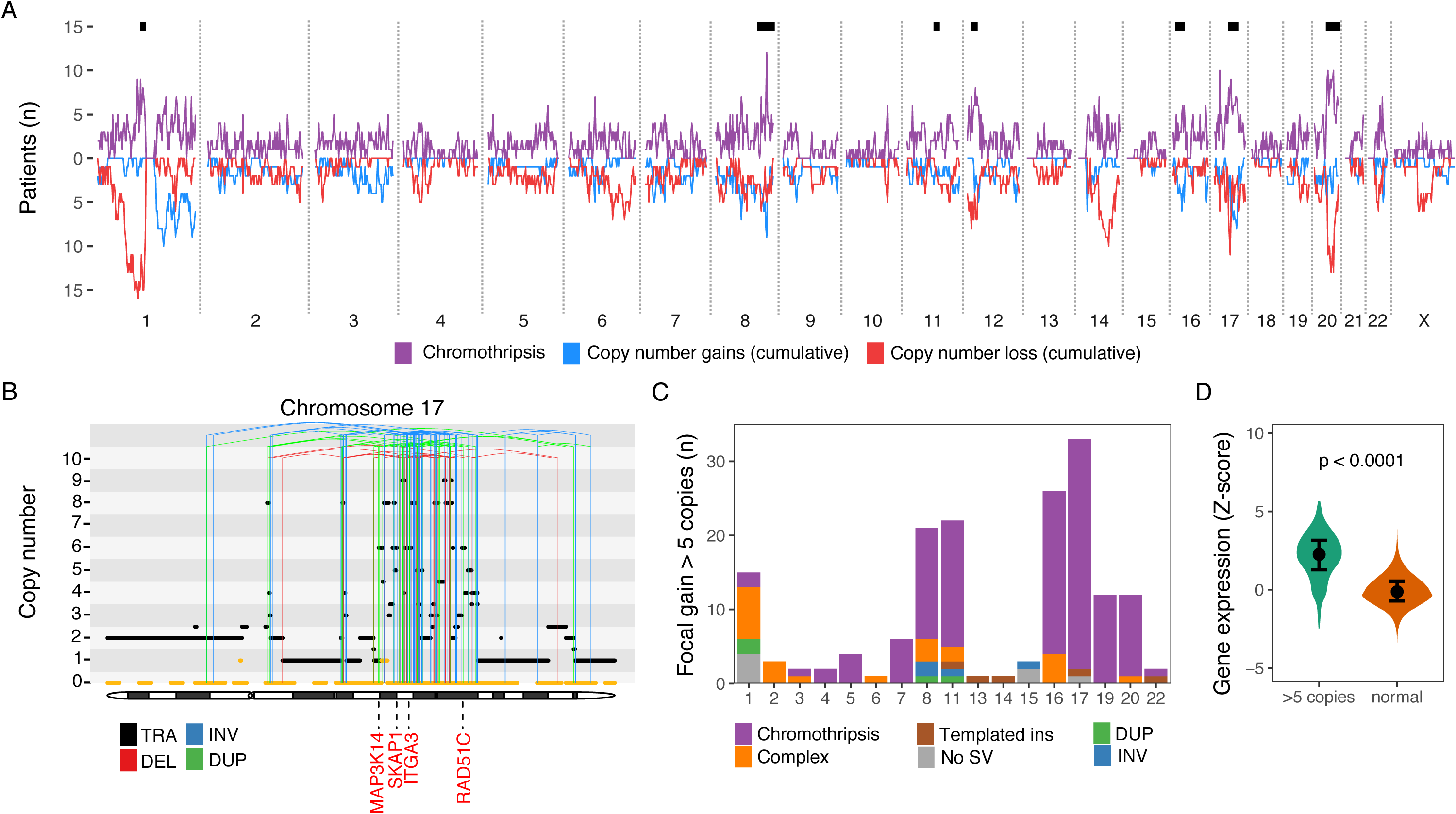
Wide-spread involvement of chromothripsis with gain and loss of function effects. **A)** Distribution of chromothripsis involvement across the genome (above), with associated cumulative copy number changes (below). Black lines indicate hotspots of chromothripsis. **B)** Example of chromothripsis causing high-level focal gains on chromosome 17 (sample MMRF_2330_1_BM). The horizontal black line indicates total copy number; the dashed orange line minor copy number. Vertical lines represent SV breakpoints, color-coded by SV class. Selected overexpressed genes (Z-score >2) are annotated in red, including the established multiple myeloma driver gene *MAP3K14,* and *RAD51C*, an oncogene commonly amplified in breast cancer (6 copies). **C)** Number of focal segments with >5 copies across chromosomes, colored by SV class. **D)** Violin plot showing gene expression Z-scores of genes involved by high-level gains (>5 copies) compared with expression of the same genes in patients where they have normal copy number. Points with error bars show median and interquartile range.

### Functional implications of recurrent and rare structural variants

To determine the functional implications of rare SVs, we first identified all simple and complex SVs where at least one breakpoint was potentially involved in a recurrent event. SVs were assigned to categories of recurrent events in a hierarchical fashion, at each step considering the SVs that had not yet been assigned. First, 9 % of SVs were assigned as canonical translocations, on the basis of one or more breakpoint involving an immunoglobulin locus or a canonical partner gene; 6.2 % were the cause of a known recurrent CNA; 4.7 % were involved an SV hotspot; and 17.4 % had one or more breakpoints within a wide GISTIC peak. Thirty-seven percent of all simple and complex SVs had one or more breakpoint in a recurrently involved region, leaving 63 % (n = 5577) as rare events (**Figure 8A**). Given the permissive criteria we used to define a recurrent event, this is likely to be a conservative estimate of their true contribution. There were striking differences in the proportion of rare and recurrent events between SV classes: the vast majority of single tandem duplications, deletions and inversions were rare, while > 75 % of chromothripsis, chromoplexy and templated insertions involved at least one recurrent region (**Figure 8A-B**). While this may be expected to some extent from the number of breakpoints involved by each event, it also supports the notion that complex events involve regions that are important in multiple myeloma and are therefore positively selected. Considering the individual breakpoints of each complex event, a greater proportion of chromothripsis breakpoints were rare as compared with templated insertions (**Figure 8B**). Interestingly, 29 % of complex events remained orphan.

**Figure 8:**
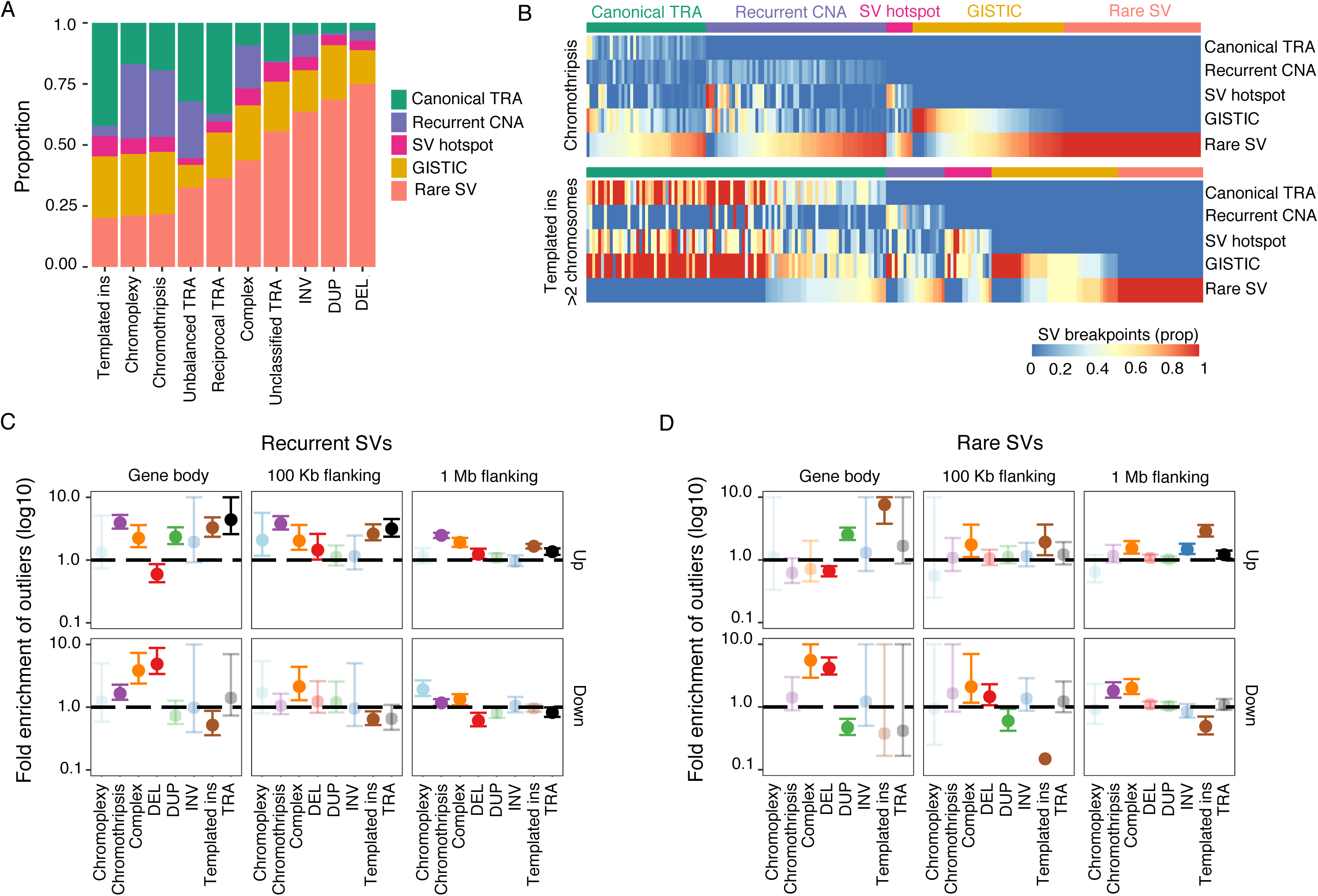
Distribution and impact of recurrent and rare structural variations. **A)** Proportion of single and complex SVs involved in one or more known or novel region of recurrent SV and/or CNA. Canonical translocation loci were prioritized (i.e. *IGH*, *IGK*, *IGL*, *MYC*, *CCND1*, *CCND2*, *CCND3*, *MAF*, *MAFA*, *MAFB* and *MMSET*), followed by known recurrent aneuploidies, SV hotspots and GISTIC peaks (See also **Tables S5-S6**). The leftover category “rare SV” constitutes those events where every breakpoint fell outside of the above-mentioned recurrently involved regions. For example, a chromothripsis event with 100 breakpoints will be classified as “Canonical TRA” if one breakpoint falls within the *IGH* locus, even if the remaining 99 breakpoints are in non-recurrent regions. **B)** Heatmaps showing the proportion of breakpoints falling into each of the categories in **A)**, displaying chromothripsis events (above) and templated insertion event involving more than two chromosomes (below). Each column in the heatmaps represent one complex event, arranged and color-coded (above the heatmap) according to the classification in **A)**. Rows in the heatmap show the proportion of breakpoints belonging to each complex event that fall into each of the categories, e.g., Canonical TRA, Recurrent CNA, etc. The sum of proportions may exceed 1 for each event, because many of the recurrent events overlap, e.g. the same region may be both recurrently translocated, an SV hotspot, and a GISTIC peak. Breakpoints falling in that region will be counted towards each of the categories. **C-D)** Enrichment analysis of outlier gene expression (Z-score +/- 2) by proximity to recurrent (**C**) and rare (**D**) SVs, showing the log10 fold-enrichment (y-axis) for each SV class (x-axis), compared with a permutation-based background model. Enrichment for overexpression outliers above and outlier reduced expression below. Columns show outlier enrichment according to the distance from SV breakpoints to involved genes.

Determining the biological implications of rare events is challenging because genomic drivers are typically defined based on recurrence above some background level. To overcome this challenge, we evaluated the impact of rare events in aggregate, asking the question whether genes near rare SVs were more or less likely to show outlier expression levels than what would be expected if all rare SVs were passenger events^55^. Outlier expression was defined for each gene as a z-score of +/- 2. Consistent with the heterogenous nature of multiple myeloma, each patient had a median of 124 genes showing outlier expression (IQR 52-436), of which a median of 3 (IQR 1-6) and 2 (IQR 1-5) outliers were within 1 Mb of recurrent and rare SV breakpoints, respectively (**Figure S8A**). Conversely, each SV was associated with a median of 1 expression outlier (IQR 1-2), similar for recurrent and rare events. Chromothripsis and templated insertions were associated with more outliers overall (**Figure S8B-C**), consistent with the complexity of these events, sometimes associated with dramatic effects on gene expression (as shown in **Figure 7D**). Structural variants showed striking enrichment for outlier over- and under-expression of nearby genes when compared with a permutation background model (**Methods**). Recurrent and rare events showed similar overall trends, although the magnitude of effect was greater for recurrent events (**Figure 8C-D**, respectively). The directions of gene expression effects were consistent with the mechanism of each SV class. Tandem duplications and templated insertions were strongly enriched for outlier overexpression, with fewer nearby genes downregulated than expected by chance.

Tandem duplications exerted their effects only when involving the gene body; while templated insertions maintained their effect up to 1 Mb away from the gene, indicating a major effect on gene regulatory mechanisms, e.g., super-enhancer hijacking. Chromothripsis and unspecified complex events were associated with both up- and down-regulation, through direct gene involvement as well as regulatory effects.

Based on the net excess of observed outliers over the background, we could estimate the number of outliers likely caused by SVs (**Figure S8D**). Recurrent templated insertions and chromothripsis events were associated with a net excess of 1334 outliers (95 % CI 1168-1490); while recurrent events of the other classes combined were associated with an excess of 595 (95 % CI 416-759) outliers. For rare SVs, the corresponding numbers were 197 (95 % CI 155-233) and 427 (95 % CI 299-547), respectively. These numbers certainly underestimate the overall impact of SVs, as we only considered direct effects on gene expression, applying a strict definition of outlier expression. Although the specific target genes often remain elusive, these data demonstrate that rare SVs strongly influence gene expression, which may contribute to the biology of individual tumors.

## Discussion

We describe the first comprehensive analysis of SVs in a large series of multiple myeloma patients with paired whole genome and RNA sequencing. Our findings reveal how simple and complex SVs shape the driver landscape of multiple myeloma, with events ranging from common CNAs and canonical translocations to a large number of non-recurrent SVs seemingly able to exert significant biological effects. The results focus attention on the importance of SVs in multiple myeloma and on the use of whole genome analyses in order to fully understand its driver landscape.

Previous studies of SVs in multiple myeloma have focused on translocations without consideration of complex events^14,38^, and our previous WGS study of 30 patients lacked the power to perform comprehensive driver discovery^20^. Here, applying a robust statistical approach^37^, we identified 100 SV hotspots, 85 of which have not previously been reported. Integrated analysis of copy number changes, gene expression and the distribution of SV breakpoints revealed 31 new potential driver genes, including the emerging drug targets *TNFRSF17* (BCMA) and *SLAMF7*^41,56^. Overall, our comprehensive catalogue of SV hotspots expands the current understanding of driver gene deregulation by recurrent CNAs and SVs and identified promising new driver candidates.

From a pan-cancer perspective, the SV landscape of multiple myeloma is characterized by a lower SV burden and less genomic complexity than in many solid tumors^2,6,11^. For example, we did not find any classical double minute chromosomes with tens to hundreds of amplified copies, nor did we find any of the recently proposed complex SV classes *pyrgo*, *rigma* and *tyfonas*^6^. Nonetheless, we found that complex SVs play a crucial role in shaping the genome of multiple myeloma patients. There are 3 major types of complex events seen in multiple myeloma, each of which is characterized by distinct junctional features and distribution patterns. Chromothripsis breakpoints are distributed across the genome in a chaotic fashion and are a common cause of recurrent deletions and gains involving tumor suppressor genes and oncogenes. While chromoplexy accounts for only a small proportion of SV breakpoints it emerged as an important cause of chained events associated with deletions potentially inactivating a range of tumor suppressor genes distributed across multiple chromosomes. In contrast, templated insertions result in amplification and concatenation of multiple SV hotspots, involving oncogenes and/or super-enhancers that may result in gene overexpression. The frequency of templated insertions emphasizes the importance of enhancer hijacking in the pathogenesis of multiple myeloma and the relevance of transcriptional control in respect to both its biology and therapy^2,14^.

A common feature of these 3 classes of SVs is the simultaneously deregulation of multiple driver genes as part of a single event. Such multi-driver events are of particular importance in myeloma progression as they can provide an explanation for the rapid changes in clinical behavior that are frequently seen in the clinic^57^. Such changes may be the result of “punctuated evolution” that occurs as a result of catastrophic intracellular events leading to the complex SVs we describe.

Gene deregulation by SV is a major contributor to the biology of multiple myeloma constituting a hallmark feature of its genome. However, the majority of SVs do not involve known drivers, SV hotspots or regions of recurrent CNV, implying they may simply be passenger variants irrelevant to tumor biology. However, we show a remarkable enrichment for gene expression outliers in proximity to rare SVs, suggesting that rare SVs may make a significant contribution to the biological diversity of multiple myeloma, including the mechanisms of disease progression and drug resistance.

## Methods

### Patients and samples

As a discovery cohort, we analyzed data from 762 patients with newly diagnosed multiple myeloma enrolled in the CoMMpass study (NCT01454297). As a validation cohort, we combined two multiple myeloma WGS datasets with a total of 52 patients^20,27,58^.

### Whole genome sequencing and analysis

CoMMpass samples underwent low coverage long-insert WGS (median 4-8X). Paired-end reads were aligned to the human reference genome (HRCh37) using the Burrows Wheeler Aligner, BWA (v0.7.8). Somatic variant calling was performed using Delly (v0.7.6)^5^ for SVs and tCoNuT for CNAs (https://github.com/tgen/MMRF_CoMMpass/tree/master/tCoNut_COMMPASS). The validation dataset was analyzed and comprehensively annotated as previously described^20,26^.

### RNA sequencing analysis and fusion calling

CoMMpass samples underwent RNA sequencing to a target coverage of 100 million reads. Paired-end reads were aligned to the human reference genome (HRCh37) using STAR v2.3.1z^59^. Transcript per million (TPM) gene expression values were obtained using Salmon v7.2^60^.

### Classification of structural variants

Each pair of structural variant breakpoints (i.e. deletion, tandem duplication, inversion or translocation) was classified as a single event, or as part of a complex event (i.e. chromothripsis, chromoplexy or unspecified complex)^2,20^. Translocation-type events were classified as single when involving no more than two breakpoint pairs and two chromosomes, subdivided into reciprocal translocations, unbalanced translocations, templated insertion or unspecified translocation as previously described^2,20^. Templated insertions could be either simple or complex, depending on the number of breakpoints and chromosomes involved, but was always defined by translocations associated with copy number gain. Chromothripsis was defined by more than 10 interconnected SV breakpoint pairs associated with oscillating copy number across one or more chromosomes. The thresholds of 10 breakpoints was imposed as a stringent criterion to avoid overestimating the prevalence of chromothripsis. Chromoplexy was defined by <10 interconnected SV breakpoints across >2 chromosomes associated with copy number loss. Patterns of three or more interconnected breakpoint pairs that did not fall into either of the above categories were classified as unspecified “complex”^20^.

### Mutational signature analysis

SNV calls from whole exome sequencing were subjected to mutational signature fitting, using the previously described R package *mmsig*^26^. High APOBEC mutational burden was defined by an absolute contribution of APOBEC mutations (mutational signatures 2 and 13) in the 4^th^ quartile among patients with evidence of APOBEC activity^26^.

### Structural basis for recurrent CNAs in multiple myeloma

We applied the following workflow to determine the structural basis for each recurrent CNA in multiple myeloma (**Table S2**). First, we identified in each patient every genomic segment involved by recurrent copy number gain or loss. Gains were defined by total copy number (CN) >2; loss as a minor CN = 0. Second, we called whole arm events in cases where the CN was equal to the 2 Mb closest to the centromere and telomere, and >90 % of the arm had the same CN state. Third, for segments that did not involve the whole arm, we searched for SV breakpoints responsible for the CNA within 50 kb of the CN segment ends. Finally, and intrachromosomal CNAs without SV support were classified as unknown. We applied Fisher’s exact test to compare the proportion of CNAs explained by an SV for each of the recurrent events, requiring Bonferroni-Holm adjusted p-values < 0.05 for statistical significance.

Multi-driver events were defined by two or more independent driver copy number segments and/or canonical driver translocations caused by the same SV (simple or complex).

### Copy number changes associated with structural variant breakpoints

To determine the genome-wide footprint of copy number changes resulting from SVs, we employed an “SV-centric” workflow, as opposed to the CNA-centric workflow described above. For each SV breakpoint, we searched for a change in copy number within 50 kb. If more than one CNA was identified, we selected the shortest segment. Deletion and amplification CNAs were defined as changes from the baseline of that chromosomal arm. As a baseline, we considered the average copy number of the two Mb closest to the telomere and centromere, respectively. This is important because deletions are often preceded by large gains, particularly in patients with HRD. In those cases, we are interested in the relative change caused by deletion, not the total CN of that segment (which may still be ≥ 2). We estimated the proportion of breakpoints associated with copy number gain or loss across patients, collapsing the data in 2 Mb bins across the genome. Confidence intervals were estimated using bootstrapping and the quantile method. For the purposes of plotting (**Figures 4A**, **6A** and **7A**), we divided the SV-associated CNAs into bins of 2 Mb. The resulting cumulative CNA plot shows the number of patients with an SV-associated deletion or amplification.

### Hotspots of structural variation breakpoints

To identify regions enriched for SV breakpoints, we employed the statistical framework of piecewise constant fitting (PCF). In principle, the PCF algorithm identifies regions where SVs are positively selected, based on enrichment of breakpoints with short inter-breakpoint distance compared to the expected background. We used the computational workflow previously described by^37^. In brief, negative binomial regression was applied to model local SV breakpoint rates under the null hypothesis (i.e. absence of selection), taking into account local features such as gene expression, replication time, non-mapping bases and histone modifications. The PCF algorithm can define hotspots without the use of binning, based on a user-defined smoothing parameter and threshold of fold-enrichment compared to the background. This allows for hotspots of widely different sized to be identified, depending on the underlying biological processes. To avoid calling hotspots driven by highly clustered breakpoints in a few samples, we also set a minimum threshold of 8 samples involved (∼ 1 % of the cohort) to be considered hotspot, as previously reported^37^. Despite this threshold, we found that complex SVs with tens to hundreds of breakpoints in a localized cluster (particularly chromothripsis) came to dominate the results. To account for this, we ran the PCF algorithm twice on different subsets of the data: 1) considering all breakpoints of non-clustered SVs (simple classes and templated insertions); and 2) including all SV classes, but randomly downsampling the data to include only one breakpoint per 500 kb per patient. Final output from both approaches was merged for downstream analysis.

The full SV hotspot analysis workflow is attached as **Data S1**, drawing on generic analysis tools that we have made available on github (https://github.com/evenrus/hotspots/tree/hotornot-mm). The workflow is easily portable to other datasets and includes utilities for data preparation and background model generation as well as hotspot discovery and visualization.

### Functional classification of structural variation hotspots

SV hotspots were classified based on local copy number and gene expression data as gain of function, loss of function, or unknown.

Copy number data was integrated from two complimentary approaches. First, we applied the GISTIC v2.0 algorithm to identify wide peaks of enrichment for chromosomal amplification or deletion (FDR < 0.1), using standard settings^39^. Second, we considered the cumulative copy number profiles of each hotspot, considering only the patients with SV breakpoints within the region, looking for more subtle patterns of recurrent CNA that was not picked up in the genome-wide analysis.

To determine the effects of SV hotspot involvement on gene expression, we performed multivariate modeling using the *limma* package in R^61^. Gene expression count data were first normalized using the DESeq/vst package, which allows the use of linear models^62^. We went on to estimate the effect SV hotspot involvement on every expressed gene after adjustment for the primary molecular classes, *MYC* translocation and gain1q21. Significantly altered expression (Bonferroni-Holm adjusted p-values < 0.01) of a putative driver gene within 5 Mb of the hotspot was considered as supporting evidence for a driver role.

### Identification of putative driver genes involved by SV hotspots

Multiple lines of evidence were considered to identify driver genes involved by SV hotspots. Evidence of a putative driver gene included: 1) involved by driver SNVs in multiple myeloma^19,20^; 2) included in the COSMIC cancer gene census (https://cancer.sanger.ac.uk/census); 3) designated as putative driver gene in The Cancer Genome Atlas^63–66^; 4) enrichment of SV breakpoints in or around the gene; 5) nearby peak of SV-related copy number gain or loss; 6) SV classes and recurrent copy number changes corresponding to a known role of that gene in cancer (i.e. oncogene or tumor suppressor); and 7) differential gene expression. Having identified candidate driver genes involved by SV hotspots, we reviewed the literature for evidence of a role in multiple myeloma (**Table S4**).

### Histone H3K27ac and super-enhancers

Active enhancer (H3K27ac) and super-enhancer data from primary multiple myeloma cells was obtained from^40^. Enrichment of super-enhancers in hotspots was assessed using a Poisson test, comparing the super-enhancer density within 100 kb of hotspots with the remaining mappable genome.

### Enrichment of germline predisposition loci

Germline predisposition SNPs for multiple myeloma were obtained from a recent meta-analysis of GWAS-studies^48,49^. Poisson test for enrichment in hotspots was performed as for super-enhancers. We also simulated 1000 sets of hotspots of identical size using a bed shuffle implementation in R^67^, estimating the probability under the null hypothesis to obtain a more extreme result than the observed data. As a final control measure, we repeated the same analysis for loci associated with development of a completely unrelated condition. We selected schizophrenia as a control condition because the pathogenesis is distinct from multiple myeloma, with abundant data on germline risk loci^68^.

### External validation of hotspots

Hotspots identified by the PCF algorithm were validated in our external cohort of 52 patients. We applied one-sided poisson tests for enrichment of SV breakpoints within each defined hotspot region as compared with the mappable non-hotspot genome (FDR < 0.1). To avoid bias from highly clustered events, we first downsampled the data as described for the main hotspot analysis, randomly selecting a single breakpoint per 500 kb per patient.

### Chromothripsis hotspot analysis

Empirical background models showed very poor ability to predict the distribution of chromothripsis breakpoints, as may be expected if DNA breaks in chromothripsis tend to be random. To identify regions enriched for chromothripsis, we applied the PCF algorithm with a uniform background, only adjusting for non-mapping bases.

### Gene expression analysis of rare structural variants

To determine whether rare SVs are functionally relevant, we assessed their aggregated effect on gene expression outliers, adopting an approach previously published for germline SVs^55^. First, we selected all genes in our cohort requiring >0 TPM expression in >25 % of patients and a median expression level of > 1. Second, we defined expression outliers with a Z-score of +/- 2 for each gene. Genes involved by SVs was defined separately for deletion/tandem duplication type SVs and translocation/inversion types. For deletions and tandem duplications, only focal events were considered (size < 3 Mb), and genes were considered directly involved if the gene was between the breakpoints, or if a breakpoint transected the gene. Genes within 100kb or 100kb – 1 Mb of a breakpoint were considered as separate categories. For translocations and inversions, direct involvement was considered when the breakpoint transected the gene. Genes within 100 Kb and 100 Kb - 1 Mb were considered as separate categories. Having defined gene expression outliers and genes involved by SVs, we compared the observed number of outliers within each category with the numbers expected by chance. In the background permutation model, we randomly shuffled the outlier status (up, down or non-outlier) of each gene. The median frequency across permutations was used as background estimates, with 95 % confidence intervals estimated by the quantile method. Fold enrichment and net excess of outliers was based on the observed frequency divided by the background frequency and the upper and lower bounds of its 95 % confidence interval.

### Data and software availability

All the raw data used in the study are already publicly available (dbGap: phs000748.v1.p1, dbGap: phs000348.v2.p1 and European Genome phenome Archive: EGAD00001001898. Analysis was carried out in R version 3.6.1. The full analytical workflow in R to identify hotspots of structural variants is provided in **Data S1**. All other software tools used are publicly available. Analysis code will be provided by the Lead Contact upon request.

## Acknowledgements

This work is supported by the Memorial Sloan Kettering Cancer Center NCI Core Grant (P30 CA 008748), the Multiple Myeloma Research Foundation (MMRF), and the Perelman Family Foundation.

K.H.M. is supported by the Haematology Society of Australia and New Zealand New Investigator Scholarship and the Royal College of Pathologists of Australasia Mike and Carole Ralston Travelling Fellowship Award.

G.J.M is supported by The Leukemia Lymphoma Society.

## Author contributions

F.M. designed and supervised the study, collected and analyzed data and wrote the paper. E.H.R. designed the study, collected and analyzed data and wrote the paper. G.J.M. and O.L. collected and analyzed data and wrote paper. D.G., V.Y, K.M. and B.D.D, analyzed data and wrote the paper. P.J.C., E.P., G.G., D.L. and N.A., analyzed data. E.M.B., C.A., M.H., A.D., Y.Z., P.B., D.A., K.C.A., P.M., N.B., H.A.L., N.M., J.K., G.M., collected data.

## Supplementary materials

**Table S1: Cohort summary.** Baseline clinical variables and availability of sequencing data from patients in the CoMMpass cohort.

**Table S2: Recurrent CNAs in multiple myeloma**. Chromosomal coordinates used to define recurrent CNAs in multiple myeloma.

**Table S3: Proportion of CNAs explained by an SV in CoMMpass and the validation dataset.** For each of the recurrent CNAs in multiple myeloma, we report the proportion of events explained by an SV in the CoMMpass and the validation dataset. Odds ratios (OR) are from Fisher’s exact test, with Bonferroni-Holm adjusted p-values.

**Table S4: SV hotspots in multiple myeloma.** Summary table of all SV hotspots. Detailed explanations for each column are provided as a second sheet in the table excel file.

**Table S5: GISTIC amplification peaks.** Wide peaks of chromosomal gains as defined by the GISTIC algorithm (FDR < 0.1).

**Table S6: GISTIC deletion peaks.** Wide peaks of chromosomal loss as defined by the GISTIC algorithm (FDR < 0.1).

**Table S7: Evaluation of fragile sites in multiple myeloma.** Shows all genes with three or more patients harboring deletion breakpoints within the gene body. Genes are assigned as known fragile sites if their genomic coordinates intersect with previously reported fragile sites in cancer, or as potential fragile sites if their size exceeds 500 Kb and they are late replicating. Late replicating was defined here as below the mean (i.e. normalized replication time below zero) across the genome of lymphoblastoid cells.

**Table S8: Germline predisposition loci in multiple myeloma and associated SV hotspots.** Shows all previously described germline predisposition loci for multiple myeloma, including the genes involved (SNP_gene) and suspected target genes (SNP_target), as previously reported. We go on to report the overlapping SV hotspot (see also **Table S4**).

**Table S9: Validation of SV hotspots.** Number and proportion of patients with an SV in each hotspot across CoMMpass and the validation dataset, with results from Poisson-tests for hotspot enrichment in the validation dataset.

**Table S10: Hotspots of chromothripsis.** Hotspot regions for chromothripsis identified by the PCF algorithm when run exclusively with chromothripsis breakpoints.

## References

1. Zhang, Y. et al. A Pan-Cancer Compendium of Genes Deregulated by Somatic Genomic Rearrangement across More Than 1,400 Cases. Cell Rep 24, 515–527, doi:10.1016/j.celrep.2018.06.025 (2018).

2. Li, Y. et al. Patterns of structural variation in human cancer. bioRxiv, 181339, doi:10.1101/181339 (2017).

3. Yang, L. et al. Diverse Mechanisms of Somatic Structural Variations in Human Cancer Genomes. Cell 153, 919–929, doi:https://doi.org/10.1016/j.cell.2013.04.010 (2013).

4. Maciejowski, J. & Imielinski, M. Modeling cancer rearrangement landscapes. Current Opinion in Systems Biology 1, 54–61, doi:https://doi.org/10.1016/j.coisb.2016.12.005 (2017).

5. Rausch, T. et al. DELLY: structural variant discovery by integrated paired-end and split-read analysis. Bioinformatics 28, i333–i339, doi:10.1093/bioinformatics/bts378 (2012).

6. Hadi, K. et al. Novel patterns of complex structural variation revealed across thousands of cancer genome graphs. bioRxiv, 836296, doi:10.1101/836296 (2019).

7. Mitchell, T. J. et al. Timing the Landmark Events in the Evolution of Clear Cell Renal Cell Cancer: TRACERx Renal. Cell 173, 611–623.e617, doi:10.1016/j.cell.2018.02.020 (2018).

8. Glodzik, D. et al. Mutational mechanisms of amplifications revealed by analysis of clustered rearrangements in breast cancers. Annals of oncology: official journal of the European Society for Medical Oncology / ESMO 29, 2223–2231, doi:10.1093/annonc/mdy404 (2018).

9. Lee, J. J. et al. Tracing Oncogene Rearrangements in the Mutational History of Lung Adenocarcinoma. Cell 177, 1842–1857.e1821, doi:10.1016/j.cell.2019.05.013 (2019).

10. Zhang, C.-Z. et al. Chromothripsis from DNA damage in micronuclei. Nature 522, 179–184, doi:10.1038/nature14493 (2015).

11. Wala, J. A. et al. Selective and mechanistic sources of recurrent rearrangements across the cancer genome. bioRxiv, 187609, doi:10.1101/187609 (2017).

12. Maciejowski, J., Li, Y., Bosco, N., Campbell, P. J. & de Lange, T. Chromothripsis and Kataegis Induced by Telomere Crisis. Cell 163, 1641–1654, doi:10.1016/j.cell.2015.11.054 (2015).

13. Manier, S. et al. Genomic complexity of multiple myeloma and its clinical implications. Nat Rev Clin Oncol 14, 100–113, doi:10.1038/nrclinonc.2016.122 (2017).

14. Barwick, B. G. et al. Multiple myeloma immunoglobulin lambda translocations portend poor prognosis. Nature communications 10, 1911, doi:10.1038/s41467-019-09555-6 (2019).

15. Bolli, N. et al. Genomic patterns of progression in smoldering multiple myeloma. Nature communications 9, 3363, doi:10.1038/s41467-018-05058-y (2018).

16. Misund, K. et al. MYC dysregulation in the progression of multiple myeloma. Leukemia, doi:10.1038/s41375-019-0543-4 (2019).

17. Walker, B. A. et al. Translocations at 8q24 juxtapose MYC with genes that harbor superenhancers resulting in overexpression and poor prognosis in myeloma patients. Blood Cancer J 4, e191, doi:10.1038/bcj.2014.13 (2014).

18. Walker, B. A. et al. A compendium of myeloma-associated chromosomal copy number abnormalities and their prognostic value. Blood 116, e56–e65, doi:10.1182/blood-2010-04-279596 (2010).

19. Walker, B. A. et al. Identification of novel mutational drivers reveals oncogene dependencies in multiple myeloma. Blood 132, 587–597, doi:10.1182/blood-2018-03-840132 (2018).

20. Maura, F. et al. Genomic landscape and chronological reconstruction of driver events in multiple myeloma. Nature communications 10, 3835, doi:10.1038/s41467-019-11680-1 (2019).

21. Walker, B. A., et al. A high-risk, Double-Hit, group of newly diagnosed myeloma identified by genomic analysis. Leukemia 33, 159–170, doi:10.1038/s41375-018-0196-8 (2019).

22. Weinhold, N. et al. Clonal selection and double-hit events involving tumor suppressor genes underlie relapse in myeloma. Blood 128, 1735–1744, doi:10.1182/blood-2016-06-723007 (2016).

23. Jones, J. R. et al. Clonal evolution in myeloma: the impact of maintenance lenalidomide and depth of response on the genetics and sub-clonal structure of relapsed disease in uniformly treated newly diagnosed patients. Haematologica, doi:10.3324/haematol.2018.202200 (2019).

24. Korbel, J. O. & Campbell, P. J. Criteria for inference of chromothripsis in cancer genomes. Cell 152, 1226–1236, doi:10.1016/j.cell.2013.02.023 (2013).

25. Baca, S. C. et al. Punctuated evolution of prostate cancer genomes. Cell 153, 666–677, doi:10.1016/j.cell.2013.03.021 (2013).

26. Rustad, E. H. et al. Timing the Initiation of Multiple Myeloma. Sneak Peak (pre-print), doi:https://dx.doi.org/10.2139/ssrn.3409453 (2019).

27. Lohr, J. G. et al. Widespread genetic heterogeneity in multiple myeloma: implications for targeted therapy. Cancer Cell 25, 91–101, doi:10.1016/j.ccr.2013.12.015 (2014).

28. Maura, F. et al. Biological and prognostic impact of APOBEC-induced mutations in the spectrum of plasma cell dyscrasias and multiple myeloma cell lines. Leukemia 32, 1044–1048, doi:10.1038/leu.2017.345 (2018).

29. Walker, B. A. et al. APOBEC family mutational signatures are associated with poor prognosis translocations in multiple myeloma. Nature communications 6, 6997, doi:10.1038/ncomms7997 (2015).

30. Magrangeas, F., Avet-Loiseau, H., Munshi, N. C. & Minvielle, S. Chromothripsis identifies a rare and aggressive entity among newly diagnosed multiple myeloma patients. Blood 118, 675–678, doi:10.1182/blood-2011-03-344069 (2011).

31. Rausch, T. et al. Genome sequencing of pediatric medulloblastoma links catastrophic DNA rearrangements with TP53 mutations. Cell 148, 59–71, doi:10.1016/j.cell.2011.12.013 (2012).

32. Quigley, D. A. et al. Genomic Hallmarks and Structural Variation in Metastatic Prostate Cancer. Cell 174, 758–769.e759, doi:10.1016/j.cell.2018.06.039 (2018).

33. Grobner, S. N. et al. The landscape of genomic alterations across childhood cancers. Nature 555, 321–327, doi:10.1038/nature25480 (2018).

34. Nutt, S. L., Hodgkin, P. D., Tarlinton, D. M. & Corcoran, L. M. The generation of antibody-secreting plasma cells. Nat Rev Immunol 15, 160–171, doi:10.1038/nri3795 (2015).

35. Boothby, M. R., Hodges, E. & Thomas, J. W. Molecular regulation of peripheral B cells and their progeny in immunity. Genes & development 33, 26–48, doi:10.1101/gad.320192.118 (2019).

36. Mikulasova, A. et al. Microhomology-mediated end joining drives complex rearrangements and over expression of MYC and PVT1 in multiple myeloma. Haematologica, haematol.2019.217927, doi:10.3324/haematol.2019.217927 (2019).

37. Glodzik, D. et al. A somatic-mutational process recurrently duplicates germline susceptibility loci and tissue-specific super-enhancers in breast cancers. Nat Genet 49, 341–348, doi:10.1038/ng.3771 (2017).

38. Walker, B. A. et al. Characterization of IGH locus breakpoints in multiple myeloma indicates a subset of translocations appear to occur in pregerminal center B cells. Blood 121, 3413–3419, doi:10.1182/blood-2012-12-471888 (2013).

39. Mermel, C. H. et al. GISTIC2.0 facilitates sensitive and confident localization of the targets of focal somatic copy-number alteration in human cancers. Genome Biology 12, R41, doi:10.1186/gb-2011-12-4-r41 (2011).

40. Jin, Y. et al. Active enhancer and chromatin accessibility landscapes chart the regulatory network of primary multiple myeloma. Blood 131, 2138–2150, doi:10.1182/blood-2017-09-808063 (2018).

41. Cho, S. F., Anderson, K. C. & Tai, Y. T. Targeting B Cell Maturation Antigen (BCMA) in Multiple Myeloma: Potential Uses of BCMA-Based Immunotherapy. Frontiers in immunology 9, 1821, doi:10.3389/fimmu.2018.01821 (2018).

42. Affer, M. et al. Promiscuous MYC locus rearrangements hijack enhancers but mostly super-enhancers to dysregulate MYC expression in multiple myeloma. Leukemia 28, 1725–1735, doi:10.1038/leu.2014.70 (2014).

43. Glover, T. W., Wilson, T. E. & Arlt, M. F. Fragile sites in cancer: more than meets the eye. Nature reviews. Cancer 17, 489–501, doi:10.1038/nrc.2017.52 (2017).

44. Veeriah, S. et al. The tyrosine phosphatase PTPRD is a tumor suppressor that is frequently inactivated and mutated in glioblastoma and other human cancers. Proceedings of the National Academy of Sciences 106, 9435, doi:10.1073/pnas.0900571106 (2009).

45. Hussain, T., Liu, B., Shrock, M. S., Williams, T. & Aldaz, C. M. WWOX, the FRA16D gene: A target of and a contributor to genomic instability. Genes, chromosomes & cancer 58, 324–338, doi:10.1002/gcc.22693 (2019).

46. Patel, K. et al. FAM190A deficiency creates a cell division defect. The American journal of pathology 183, 296–303, doi:10.1016/j.ajpath.2013.03.020 (2013).

47. Waters, C. E., Saldivar, J. C., Hosseini, S. A. & Huebner, K. The FHIT gene product: tumor suppressor and genome “caretaker”. Cell Mol Life Sci 71, 4577–4587, doi:10.1007/s00018-014-1722-0 (2014).

48. Went, M. et al. Identification of multiple risk loci and regulatory mechanisms influencing susceptibility to multiple myeloma. Nature communications 9, 3707, doi:10.1038/s41467-018-04989-w (2018).

49. Janz, S. et al. Germline Risk Contribution to Genomic Instability in Multiple Myeloma. Frontiers in genetics 10, doi:10.3389/fgene.2019.00424 (2019).

50. Hu, G., Wei, Y. & Kang, Y. The Multifaceted Role of MTDH/AEG-1 in Cancer Progression. Clinical Cancer Research 15, 5615, doi:10.1158/1078-0432.CCR-09-0049 (2009).

51. Cortés-Ciriano, I. et al. Comprehensive analysis of chromothripsis in 2,658 human cancers using whole-genome sequencing. bioRxiv, 333617, doi:10.1101/333617 (2018).

52. Stephens, P. J. et al. Massive Genomic Rearrangement Acquired in a Single Catastrophic Event during Cancer Development. Cell 144, 27–40, doi:10.1016/j.cell.2010.11.055 (2011).

53. Barlund, M. et al. Multiple genes at 17q23 undergo amplification and overexpression in breast cancer. Cancer research 60, 5340–5344 (2000).

54. Bahrami, B. F., Ataie-Kachoie, P., Pourgholami, M. H. & Morris, D. L. p70 Ribosomal protein S6 kinase (Rps6kb1): an update. Journal of clinical pathology 67, 1019–1025, doi:10.1136/jclinpath-2014-202560 (2014).

55. Chiang, C. et al. The impact of structural variation on human gene expression. Nature Genetics 49, 692, doi:10.1038/ng.3834 (2017).

56. Campbell, K. S., Cohen, A. D. & Pazina, T. Mechanisms of NK Cell Activation and Clinical Activity of the Therapeutic SLAMF7 Antibody, Elotuzumab in Multiple Myeloma. Frontiers in immunology 9, 2551, doi:10.3389/fimmu.2018.02551 (2018).

57. Maura, F. et al. Moving From Cancer Burden to Cancer Genomics for Smoldering Myeloma: A Review. JAMA oncology, doi:10.1001/jamaoncol.2019.4659 (2019).

58. Chapman, M. A. et al. Initial genome sequencing and analysis of multiple myeloma. Nature 471, 467–472, doi:10.1038/nature09837 (2011).

59. Dobin, A. et al. STAR: ultrafast universal RNA-seq aligner. Bioinformatics 29, 15–21, doi:10.1093/bioinformatics/bts635 (2013).

60. Patro, R., Duggal, G., Love, M. I., Irizarry, R. A. & Kingsford, C. Salmon provides fast and bias-aware quantification of transcript expression. Nature methods 14, 417–419, doi:10.1038/nmeth.4197 (2017).

61. Gerstung, M. et al. Combining gene mutation with gene expression data improves outcome prediction in myelodysplastic syndromes. Nature communications 6, 5901, doi:10.1038/ncomms6901 (2015).

62. Love, M. I., Huber, W. & Anders, S. Moderated estimation of fold change and dispersion for RNA-seq data with DESeq2. Genome Biol 15, 550, doi:10.1186/s13059-014-0550-8 (2014).

63. Sanchez-Vega, F. et al. Oncogenic Signaling Pathways in The Cancer Genome Atlas. Cell 173, 321–337.e310, doi:10.1016/j.cell.2018.03.035 (2018).

64. Seiler, M. et al. Somatic Mutational Landscape of Splicing Factor Genes and Their Functional Consequences across 33 Cancer Types. Cell Reports 23, 282–296.e284, doi:https://doi.org/10.1016/j.celrep.2018.01.088 (2018).

65. Ge, Z. et al. Integrated Genomic Analysis of the Ubiquitin Pathway across Cancer Types. Cell Reports 23, 213–226.e213, doi:https://doi.org/10.1016/j.celrep.2018.03.047 (2018).

66. Knijnenburg, T. A. et al. Genomic and Molecular Landscape of DNA Damage Repair Deficiency across The Cancer Genome Atlas. Cell Reports 23, 239–254.e236, doi:https://doi.org/10.1016/j.celrep.2018.03.076 (2018).

67. Riemondy, K. A. et al. valr: Reproducible genome interval analysis in R. F1000Research 6, 1025–1025, doi:10.12688/f1000research.11997.1 (2017).

68. Huo, Y., Li, S., Liu, J., Li, X. & Luo, X.-J. Functional genomics reveal gene regulatory mechanisms underlying schizophrenia risk. Nature communications 10, 670, doi:10.1038/s41467-019-08666-4 (2019).

