## Supplementary materials for "Revealing the impact of recurrent and rare structural variants in multiple myeloma"

### Supplementary Figures

**Figure S1: Structural variants landscape in multiple myeloma. A)** SV burden and classification in our discovery (CoMMpass; left panels; n = 762) and validation (right panels; n = 52) datasets. Relative (top) and absolute (bottom) contribution of simple and complex classes (color) for each of the four SV types (bars): deletion (DEL), tandem duplication (DUP), inversion (INV) and translocation (TRA). **B**) Correlation matrix showing the breakpoint density of each SV class in the CoMMpass dataset (y-axis) and genomic features across the genome (x-axis), divided in 500 kb bins. Color and size of points are determined by the magnitude of positive (blue) and negative (red) spearman correlation coefficients. **C)** Correlation between the number of SVs of each class across patients in the CoMMpass dataset. Positive correlation between a pair of SV classes indicates that patients who have a high frequency of one SV class tends to have higher frequency of the other as well.

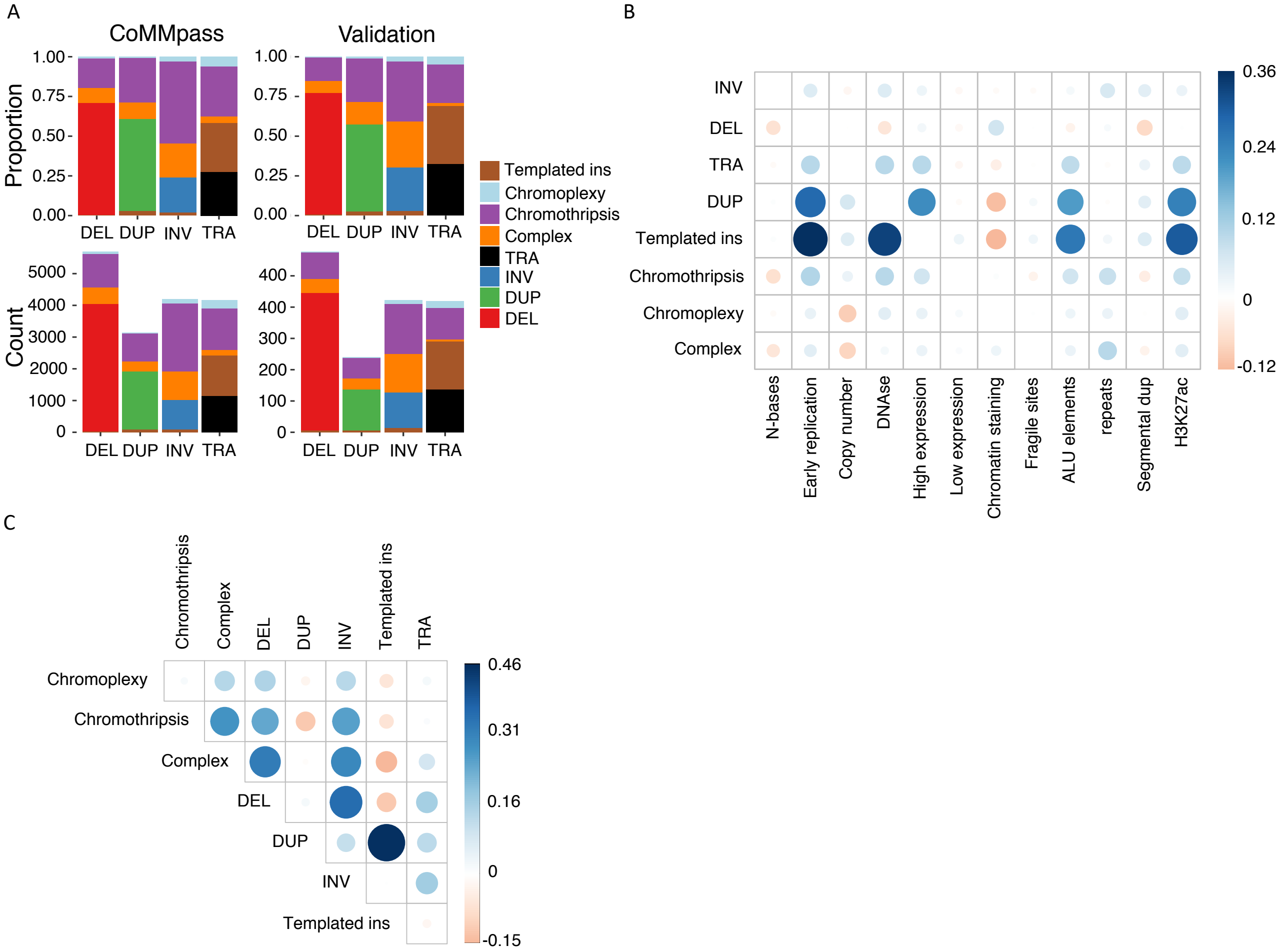

**Figure S2: Co-occurrence and mutual exclusivity of complex SV classes and key molecular features of multiple myeloma.** Heatmap of co-occurrence and mutual exclusivity of complex SV classes, disease-initiating events (e.g., HRD or *IGH*-translocations), *MYC* translocations, recurrent copy number alterations, and high APOBEC mutational burden. Color intensity indicates the magnitude of co-occurrence (green) or mutual exclusivity (purple), as evaluated by Fisher's exact test. Statistical significance is indicated by black dots (FDR < 0.1) and asterisks (Bonferroni-Holm adjusted p-values < 0.05).

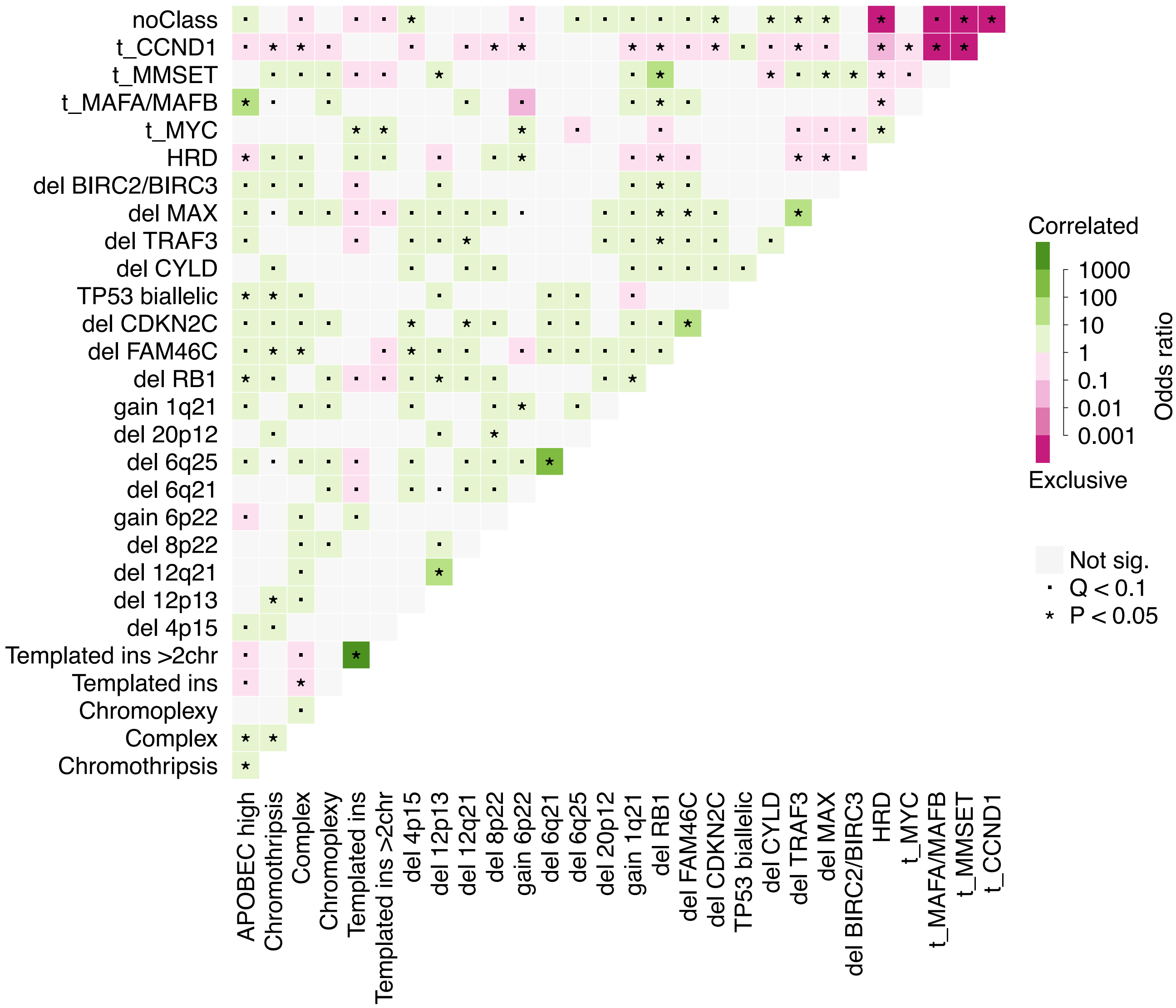

**Figure S3: Background model validation and piecewise constant fitting for hotspot discovery. A-C:** Coefficients and validation data from a negative binomial regression model to predict the background density of non-clustered SVs in genome-wide 500 kb bins (see also **Data S1**). **A)** regression coefficients (bars) +/- 1 standard error (horizontal lines). **B)** Negative binomial regression compared with random permutation, using 70 % of bins for training and 30 % for testing. Test data performance is shown here. Point estimates from each bootstrapped dataset are faded. Solid points and vertical lines represent the median and 95 % CI from bootstrapping estimates. **C)** Observed (y-axis) versus predicted (x-axis) SV breakpoint density in test data, showing p-value and rho from spearman correlation. **D)** SV breakpoints (points) across chromosome 1 (x-axis) plotted according to the log-distance from its neighboring breakpoint (y-axis). Peaks of breakpoints pointing down towards the X axis indicate hotspots. Horizontal black lines drawn in the plot area indicate hotspots as defined by the piecewise constant fitting algorithm. Breakpoints and hotspots of non-clustered SVs are shown in the upper panel; downsampling of all SV classes in the lower. **E)** Hotspots identified by two integrated approaches. **F)** Total SV burden in each patient (y-axis) and the number of hotspots identified (x-axis), colored by the presence or absence of chromothripsis. Lines with shaded background represent linear regression lines for the two groups.

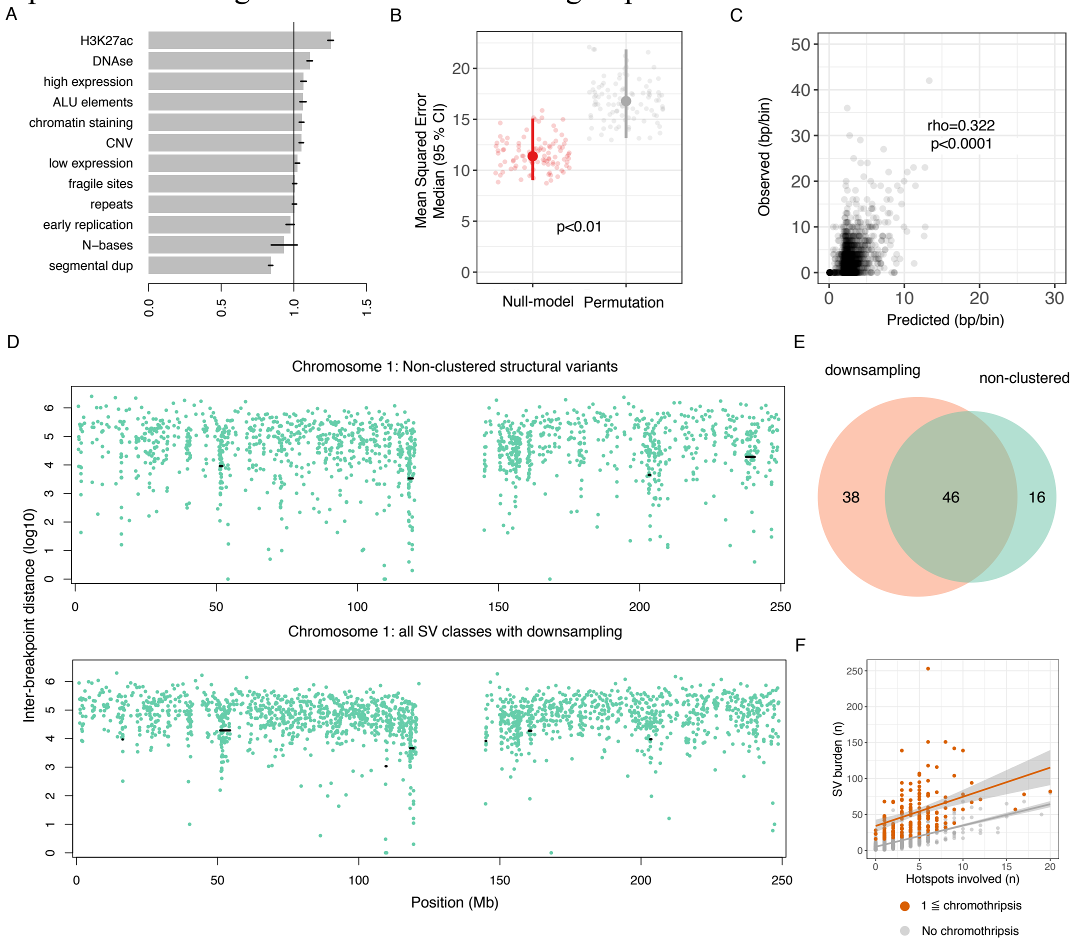

**Figure S4: Hotspots with novel putative driver genes**. Overall, we identified 31 hotspots containing novel putative driver genes. Two are shown as examples in the main manuscript (**Figure 4B-C**); the remaining 29 are displayed here. Each panel displays, from top to bottom: SV density (black), enhancer density (brown), GISTIC peaks and cumulative copy number (gain = blue, red = loss). Vertical dashed gray lines mark the loci of putative driver genes in the SV density plots. In cumulative CNA plots, the number of patients with CNA is shown on the y-axis, with chromosomal coordinates (Mb) on the x-axis. Asterisks indicate hotspots with significantly altered expression of a putative driver gene.

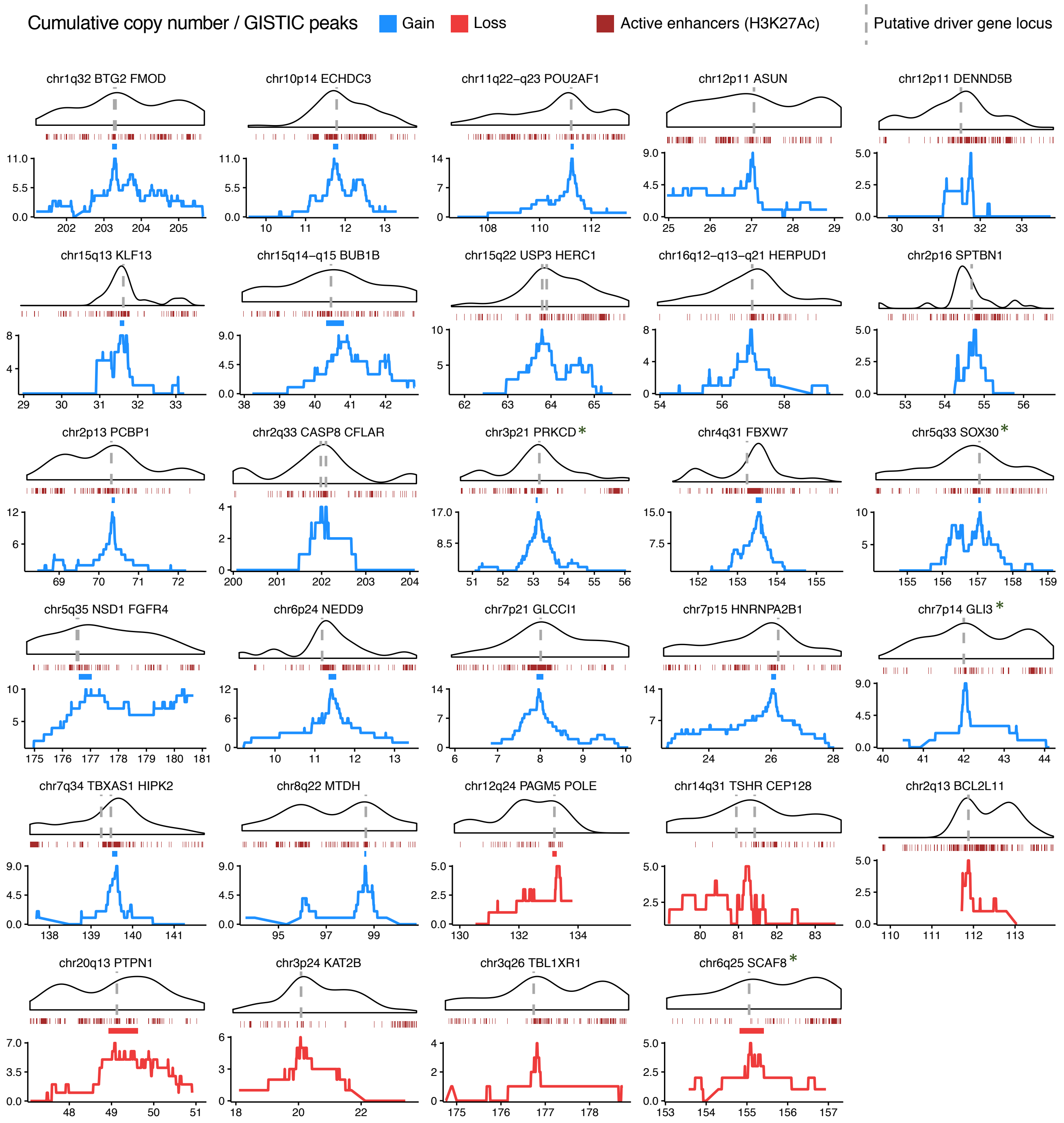

**Figure S5: Gene expression effects of SVs in gain-of-function hotspots.** Violin plots show the Z-scores of known and novel putative driver genes in gain-of-function hotspots involved by different categories of SVs, as compared with expression in the absence of SV involvement. Error bars show the median and interquartile range for each group. Canonical immunoglobulin translocation partners are shown to the left (e.g., *CCND1*, *MMSET*, *FGFR3* and *MYC*); other putative drivers to the right (e.g., *TNFRSF17*, *CD40*, *KLF2* and *KLF13*). Within each group of hotspots (canonical vs., other), we applied pairwise Wilcoxon tests, requiring Bonferroni-Holm adjusted p-values < 0.05 for statistical significance. Among canonical translocation partner genes, IG partners had significantly stronger expression effects than all other groups ($p < 0.0001$). The remaining SV categories were not different from each other but were all associated with higher expression than the absence of SV. Among other putative driver genes, a similar striking pattern was observed for translocations involving immunoglobulin genes. Moreover, non-IG translocations and tandem duplications had stronger effects than the remaining SV classes; and the presence of any SV was associated with higher expression than absence of SV. IG, immunoglobulin (i.e., *IGH*, *IGK* and *IGL*). TRA, any simple or complex translocation-type SV. DUP, single tandem duplication.

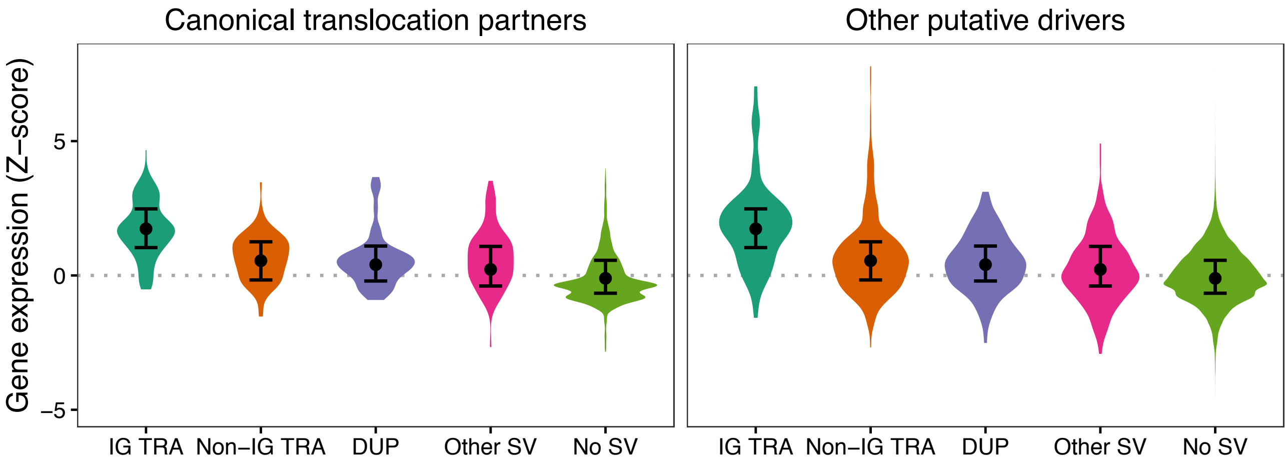

**Figure S6: Enrichment of multiple myeloma germline predisposition SNPs in SV hotspots. A-B)** Enrichment test for multiple myeloma risk loci to the left; schizophrenia risk loci as a negative control to the right (**Methods**). We used schizophrenia as a control condition to test for a general association between SV hotspots and regions prone to accumulate germline mutations. **A)** Risk loci per Mb for hotspots (HS) as compared with the remaining mappable genome (Bg). RR: rate ratio; p-values are from two-sided Poisson tests. **B)** Bar plots showing the number of risk loci within 1000 sets of simulated hotspots. Red vertical lines indicate the observed numbers. P-values were estimated by the proportion of simulations with equal or higher number of risk loci than the observed value.

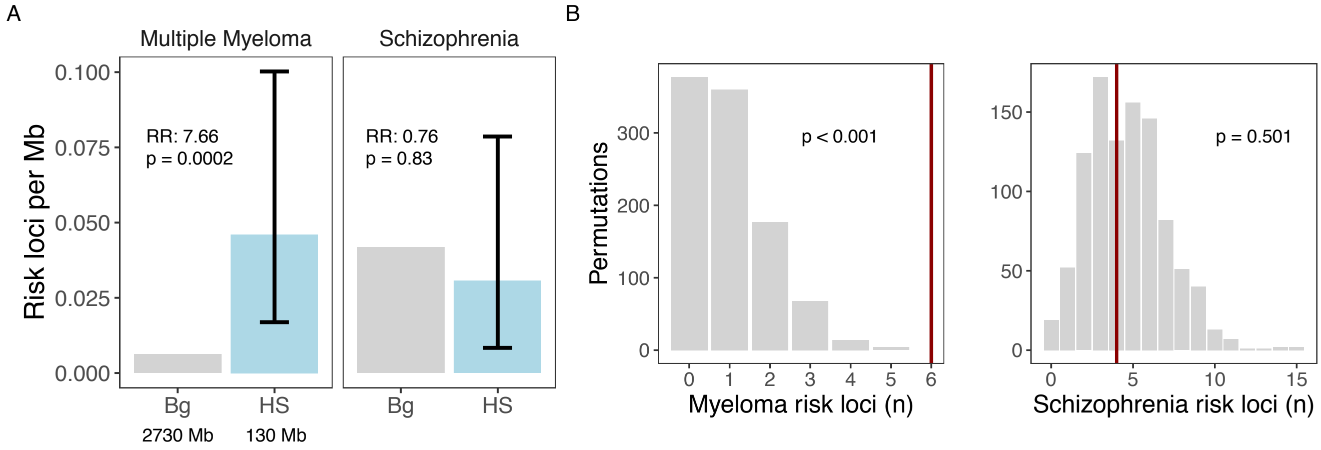

**Figure S7: Validation of SV hotspots and chromothripsis causing high-level gains. A)** Panels show the number of patients with an SV in 500 Kb bins across the genome for the CoMMpass (above) and validation (below) datasets. Dots indicate hotspots, colored by statistical significance in the validation dataset by a one-sided Poisson test. **B)** The highest number of copy number gains associated with chromothripsis in the validation dataset was 7 copies. Here displaying chromosome 8 from patient PD26419c. Copy number (y-axis) along the chromosome is shown on the x-axis: total in black and the minor allele as a dashed orange line. Colored vertical lines represent SV breakpoints; translocations (black) are annotated with the partner chromosome number. In this case all translocations were with chromosome 15. **C)** Chromothripsis with high-level gain of *RPS6KB1* (red) in sample MMRF_2814_1_BM.

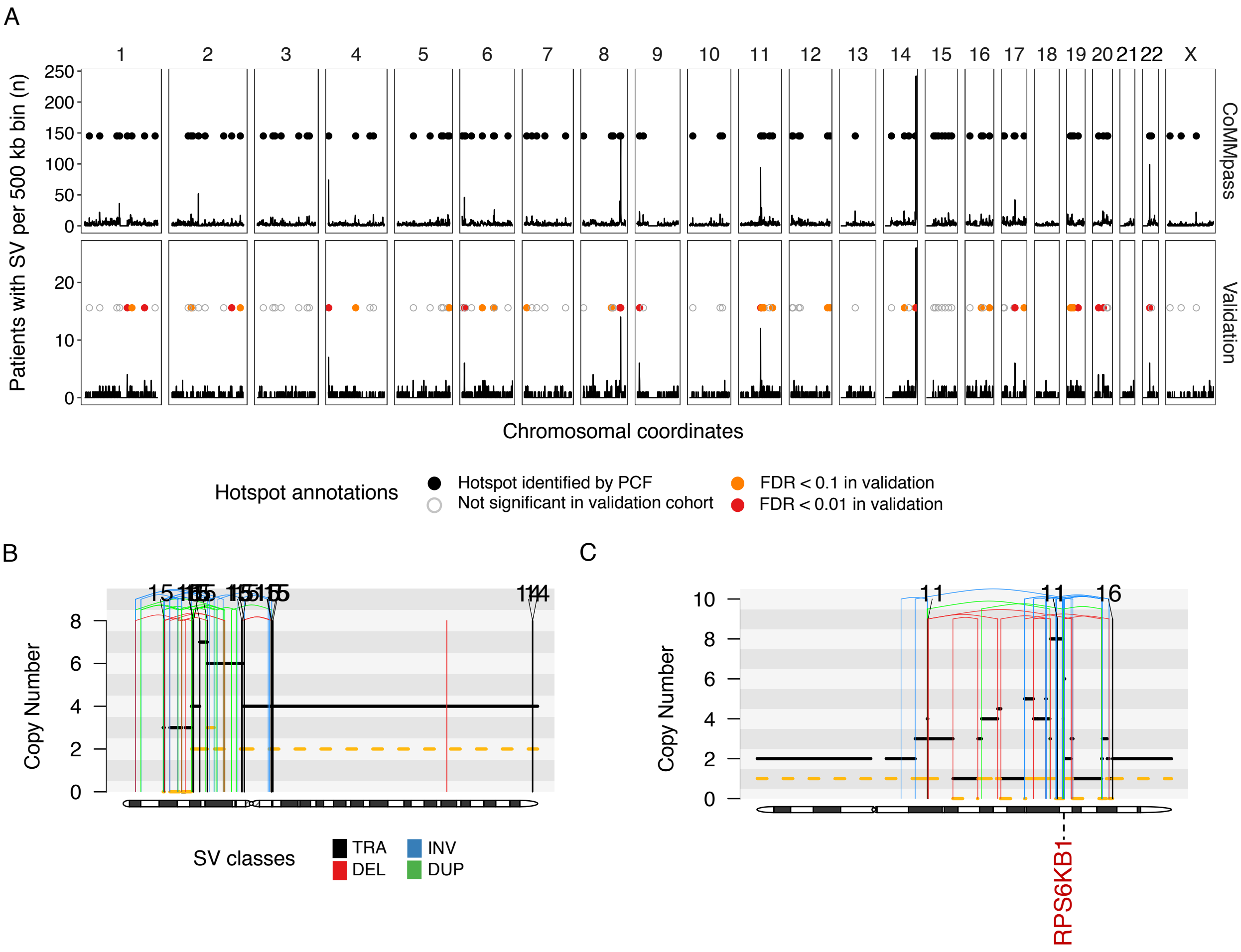

**Figure S8: Association between SVs and outlier gene expression. A)** Stacked histograms showing the number of genes in each patient that show outlier expression (Z-score of +/- 2): reduced expression to the left, overexpression to the right. Three outlier counts are shown for each patient (colors): outliers near a recurrent SV (purple), outliers near a rare SV (orange) and outliers with no SV breakpoint within 1 Mb. These counts of SV-associated outliers were compared with a random permutation model to test for enrichment of outliers near SVs, as presented in the main manuscript (**Figure 8C-D**). **B-C)** Here, we show the number of outliers associated with each individual recurrent (**B**) or rare (**C**) SV, as opposed to the patient-level summary shown in **A**). Under- and over-expression outliers are shown in the upper and lower rows, respectively; columns from left to right show the position of the SV relative to the gene. **D)** To estimate the number of gene expression outliers caused by a recurrent (purple) or rare (orange) SV, we calculated the net excess of outliers (y-axis) above the permutation-based background model. Net excess of overexpression outliers is shown to the right, reduced expression to the left. Given the potential for chromothripsis and templated insertions to involve multiple outliers in the same event (see panels **B-C**), we display the net excess for these events separately.

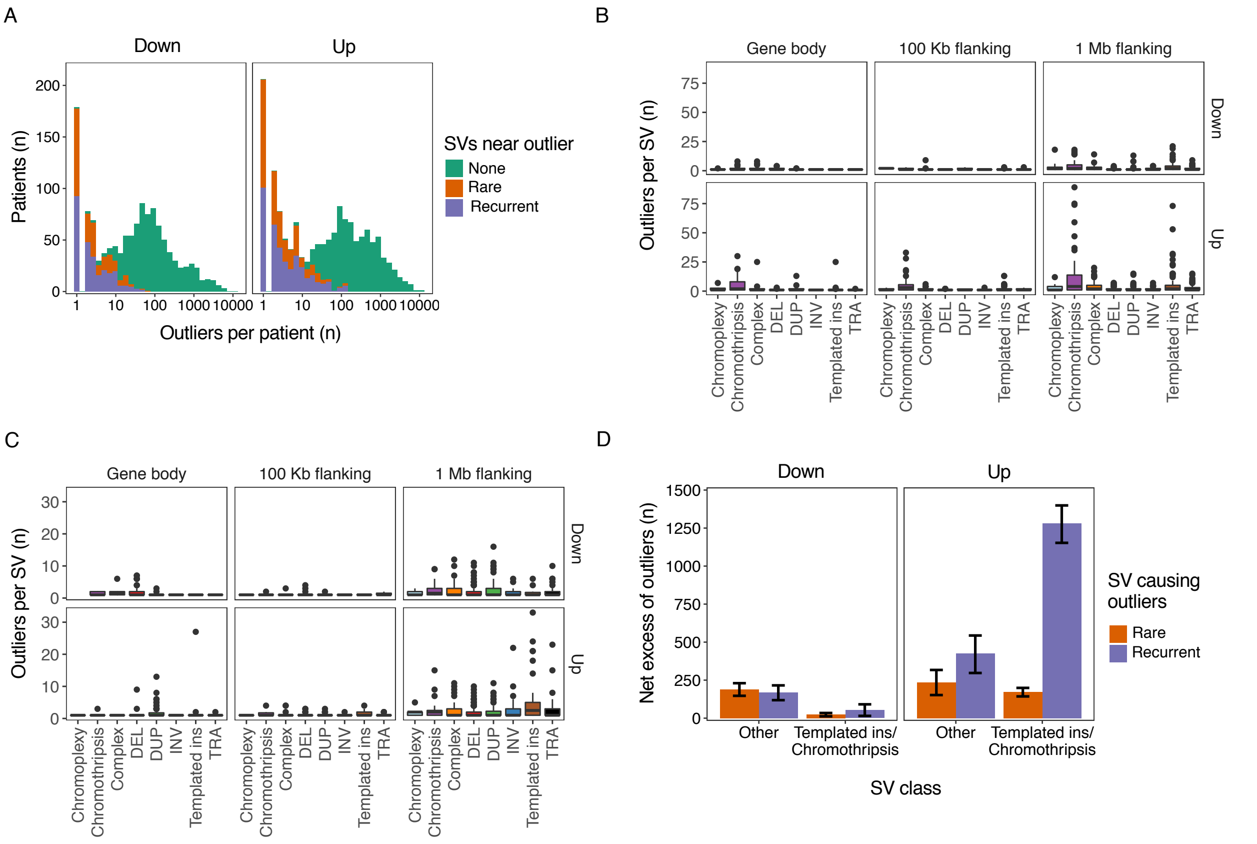
